# Modulation of cannabinoid receptor 2 alters neuroinflammation and reduces formation of alpha-synuclein aggregates in a rat model of nigral synucleinopathy

**DOI:** 10.1101/2023.08.25.554814

**Authors:** Valerie Joers, Benjamin C Murray, Caroline McLaughlin, Danielle Oliver, Hannah Staley, Jazmyn Coronado, Cindy Achat-Mendes, Sanam Golshani, Sean D. Kelly, Matthew Goodson, Danica Lee, Fredric P. Manfredsson, Bob M. Moore, Malú Gámez Tansey

## Abstract

Research into the disequilibrium of microglial phenotypes has become an area of intense focus in neurodegenerative disease as a potential mechanism that contributes to chronic neuroinflammation and neuronal loss in Parkinson’s disease (PD). There is growing evidence that neuroinflammation accompanies and may promote progression of alpha-synuclein (Asyn)-induced nigral dopaminergic (DA) degeneration. From a therapeutic perspective, development of immunomodulatory strategies that dampen overproduction of pro-inflammatory cytokines from chronically activated immune cells and induce a pro-phagocytic phenotype is expected to promote Asyn removal and protect vulnerable neurons. Cannabinoid receptor-2 (CB2) is highly expressed on activated microglia and peripheral immune cells, is upregulated in the substantia nigra of individuals with PD and in mouse models of nigral degeneration. Furthermore, modulation of CB2 protects against rotenone-induced nigral degeneration; however, CB2 has not been pharmacologically and selectively targeted in an Asyn model of PD. Here, we report that 7 weeks of peripheral administration of CB2 inverse agonist SMM-189 reduced phosphorylated (pSer129) alpha-synuclein in the substantia nigra compared to vehicle treatment. Additionally, SMM-189 delayed Asyn-induced immune cell infiltration into the brain as determined by flow cytometry, increased CD68 protein expression, and elevated wound-healing-immune-mediator gene expression. Additionally, peripheral immune cells increased wound-healing non-classical monocytes and decreased pro-inflammatory classical monocytes. *In vitro* analysis of RAW264.7 macrophages treated with lipopolysaccharide (LPS) and SMM-189 revealed increased phagocytosis as measured by the uptake of fluorescence of pHrodo *E. coli* bioparticles. Together, results suggest that targeting CB2 with SMM-189 skews immune cell function toward a phagocytic phenotype and reduces toxic aggregated species of Asyn. Our novel findings demonstrate that CB2 may be a target to modulate inflammatory and immune responses in proteinopathies.

## Background

Research into the disequilibrium of microglial phenotypes has become an area of intense focus in neurodegenerative disease as a potential mechanism that contributes to disease pathophysiology, including chronic neuroinflammation and neuronal loss (1, 2). More specifically, microglia dynamically transition from homeostatic to disease-associated phenotypes with varying transcriptional and functional profiles. In Parkinson’s disease (PD), activated microglia have been reported in postmortem brain histopathological analyses and single-cell transcriptomic studies have identified disease-specific increases in midbrain microglia clusters involved in the inflammatory response (3–6). Findings from animal models demonstrate that overexpression or delivery of fibrillar alpha-synuclein (Asyn) in the brain influences microglial phenotypes and infiltration of peripheral immune cells (5, 7–10). From a therapeutic perspective, the development of immunomodulatory strategies that dampen overproduction of pro-inflammatory cytokines from chronically activated immune cells and promote a pro-phagocytic phenotype is predicted to protect vulnerable neurons and promote toxic Asyn removal.

The cannabinoid system has been recently implicated in PD where homozygous loss-of-function diacylglycerol-lipase beta (DAGLB) mutations were linked to early-onset PD in different families of Chinese descent (11). DAGLB synthesizes 2-Arachidonoylglycerol (2-AG), which is a potent brain endocannabinoid that has high potency for cannabinoid receptors 1 and 2. Cannabinoid receptor-2 (CB2) is highly expressed on activated microglia and peripheral immune cells including circulating monocytes (12). Due to its abundant presence on immune cells, CB2 has emerged as a immunomodulatory target as it regulates various inflammatory processes including cytokine production, immune cell migration and proliferation. When evaluating the CB2 receptor expression in PD, it has been reported to be upregulated in the substantia nigra of PD patients by immunohistochemical (13, 14) and transcriptional outcome measures (15). Similarly, it is increased in mouse models of nigral degeneration (16, 17). Evidence from several show that overexpressing CB2 or pharmacologically targeting the receptor in neurotoxin animal models of PD results in dampened inflammation and in some cases neuroprotection (18, 19). Non-selective cannabinoid receptor agonists also ameliorate neurotoxin-induced neuroinflammation and neurodegeneration in PD animal models (16, 20). Yet, only one report, using genetic deletion approaches, has evaluated the role of CB2 on the clearance of Asyn aggregates in brain (21), and there has not yet been a study evaluating CB2-selective ligands on Asyn burden *in vivo*.

To further understand the role of CB2 in Asyn clearance, we sought to determine the effects of peripheral treatment with the novel CB2 inverse agonist, SMM-189, on Asyn-induced pathologies in a rat model of viral vector-mediated Asyn overexpression. Previous studies using SMM-189 demonstrated immunomodulatory effects that improve acute neuronal injury and behavioral outcomes in TBI models potentially explained by a mechanism where SMM-189 suppresses pro-inflammatory markers and increases anti-inflammatory effects (22–24). Inverse agonists bind constitutively-active CB2 receptors and lock them into an inactive conformation state preventing coupling to G_αi/o_ with a concomitant increase in cyclic AMP (cAMP) production (25). SMM-189 acts on this canonical pathway in microglia as evidenced by increased levels of nuclear phosphorylated cAMP response-element binding protein (pCREB), triggering downstream transcription of wound-healing cytokines (26). Therefore, we hypothesize that targeting CB2 will promote clearance of Asyn by altering the central inflammatory environment. In this study, we provide evidence that pharmacological inverse agonism of CB2 alters the inflammatory state of peripheral and central immune cells and reduces levels of phosphorylated human Asyn in the brain.

## Methods

### Animals and Experimental Design

Sprague Dawley male rats (16-18 weeks of age, 400-525g) were purchased from *Charles River Laboratories* and acclimated to Emory University animal facilities. Rats were pair housed in standard plexiglass cages under normal 12-hr light cycles and handled in accordance with protocols approved by the IACUC of Emory University in Atlanta, GA. Rats were administered unilateral injections of AAV2/5-human wild-type Asyn at titers of 2.9 – 4.3×10^11^ vector genomes (vg)/mL (termed “low cohort”), which was previously shown to not induce nigral neurotoxicity (27). Thereby, generating a model with overexpression of human wild-type Asyn and Asyn-mediated neuroinflammation. One week after surgery, animals were randomly selected to receive daily systemic injections of either SMM-189 (i.p., 6 mg/kg, n=12) or vehicle (i.p., n=12) and followed for 8 weeks (**Figure 1**; 2 rats per group were removed due to poor nigral targeting). The dose of SMM-189 was based on a pharmacokinetic study in mice determining therapeutic brain-blood ratios as described below. At sacrifice, animals were injected with euthasol (i.p., 150 mg/kg) and cardiac blood collected to isolate PBMCs for inflammatory gene expression using qPCR analysis. After blood collection, animals were transcardially perfused with ice-cold PBS and the frontal cortex frozen for gene analysis, the striatum frozen for western blotting analysis, and the midbrain post-fixated in 4% PFA to be processed for immunohistochemistry. A second low cohort of rats (SMM-189 n=6 or vehicle n=3 with 2 removed due to poor nigral targeting) completed the study and provided extra tissue to perform additional immunohistochemical and western blotting analyses. Plasma was also collected at baseline, 4 and 8 weeks post-AAV-hAsyn injections and CSF collected prior to perfusion to measure levels of SMM-189.

**Figure 1.**
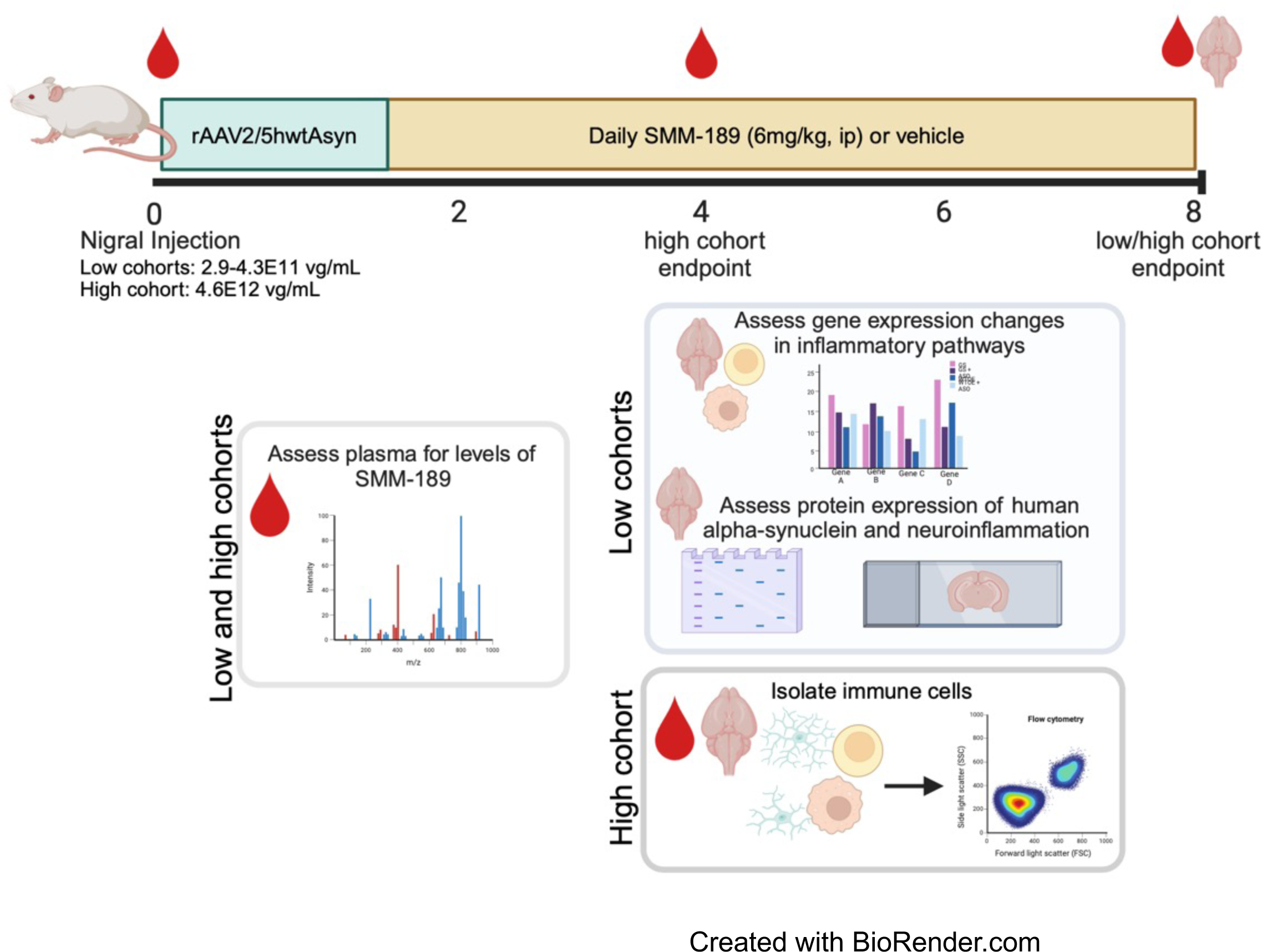
Experimental design and outcome measures. Rats were injected unilaterally with AAV2/5-human wild-type alpha-synuclein (low cohort: 2.9 – 4.3E11 vg/mL) using stereotaxic coordinates to target the substantia nigra and one week later were dosed daily with peripheral injections of SMM-189, a novel CB2 inverse agonist (6mg/kg, ip) or vehicle for 7 remaining weeks until perfusion. Blood was collected at baseline and every 4 weeks to measure plasma drug levels. Endpoint brain and PBMCs were evaluated for changes in inflammatory pathway gene expression using qPCR and the midbrain was subjected to immunohistochemistry to evaluate synuclein pathology and immune cells. An additional cohort of rats followed similar experimental design (high cohort: 4.6E12 vg/mL) to evaluate immune cells in blood (PBMCs) and brain by flow cytometry. Blood was collected at baseline, 4 weeks and 8 weeks to measure plasma drug levels and immunophenotype by flow cytometry and brains collected at either 4 or 8 weeks to immunophenotype by flow cytometry.

An additional cohort of rats (termed “high cohort”) was evaluated to determine effects of SMM-189 on immune cell populations in the brain, resident or infiltrating, and the periphery as measured by flow cytometry (**Figure 1**). To ensure infiltrating peripheral immune cells, rats were stereotaxically injected with a higher titer of AAV2/5-human wild-type Asyn (4.6×10^12^ vg/mL) and underwent an identical drug treatment timeline with animals sacrificed at 4 (SMM-189: n=5; vehicle: n=6) and 8 weeks (SMM-189: n= 7; vehicle: n=7). At sacrifice, animals were deeply anesthetized with euthasol (i.p., 150 mg/kg) and rapid decapitation performed using a sharp guillotine. Brains were collected at endpoints, and brain immune cells isolated and evaluated using multi-color flow cytometry. Additionally, blood was collected at baseline, 4 and 8 weeks (SMM-189: n= 13; vehicle: n=13) after stereotaxic surgery for PBMCs isolation to immunophenotype using flow cytometry and for plasma evaluation of SMM-189.

### Blood Brain Barrier Penetration of SMM-189 in mice

Analysis of the brain-to-blood ratios of SMM-189 were performed by Sai Life Sciences Limited (Telangana, India). All procedures of the study were in accordance with the guidelines provided by the Committee for the Purpose of Control and Supervision of Experiments on Animals (CPCSEA) as published in The Gazette of India, December 15, 1998. Prior approval of the Institutional Animal Ethics Committee (IAEC) was obtained before initiation of the study. In brief, 54 mice were weighed and divided as Group 1 (6 mg/kg, i.p.) and Group 2 (6 mg/kg, p.o.). Animals in Group 1 and Group 2 were administered intraperitoneally and orally with SMM-189 solution formulation at 6 mg/kg in 5% Ethanol, 5% Cremophor and 90% Normal Saline, respectively. The blood samples were collected under light isoflurane anesthesia at 0.08, 0.25, 0.5, 1, 2, 4, 8, 12 and 24 hr (i.p.) and 0.25, 0.5, 1, 2, 4, 6, 8, 12 and 24 hr (p.o.) in labeled micro centrifuge tube containing K_2_EDTA as anticoagulant. After blood collection, plasma was harvested by centrifugation and stored at −70 °C until analysis. Immediately after collection of blood, brain samples were collected from set of three mice at each time point at 0.08, 0.25, 0.5, 1, 2, 4, 8, 12 and 24 hr (Group 1; i.p.) and 0.25, 0.5, 1, 2, 4, 6, 8, 12 and 24 hr (Group 2; p.o.). Brain samples were homogenized using ice-cold phosphate buffer saline (pH-7.4) in a ratio of 2 (buffer):1(brain); and homogenates were stored below −70±10 °C until analysis. Total homogenate volume was three times the brain weight. All samples were processed for analysis by protein precipitation using acetonitrile and analyzed with fit-for-purpose LC-MS/MS method (LLOQ = 1.02 ng/mL for plasma and 6.09 ng/g for brain).

Non-compartmental-analysis module in Phoenix WinNonlin® (Version 6.3) was used to assess the pharmacokinetic parameters. Maximum concentration (Cmax) and time to reach the maximum concentration (Tmax) were the observed values. The areas under the concentration time curve (AUClast and AUCinf) were calculated by linear trapezoidal rule. The terminal elimination rate constant, k_e_ was determined by regression analysis of the linear terminal portion of the log plasma concentration-time curve. The terminal half-life (T1/2) was estimated as 0.693/k_e_.

### Nigral stereotaxic injections

Surgical procedures were performed under isoflurane anesthesia (induced with 5% and maintained at 1-2%). Rats were placed in a stereotaxic frame (Kopf Instruments) and administered two 2µl unilateral injections of AAV2/5-human wild-type Asyn (28, 29) into the left nigra at coordinates AP −5.3 mm, ML +2.0 mm, DV −7.2 mm and AP −6.0 mm, ML +2.0 mm and DV −7.2 mm using a glass capillary needle fitted to a Hamilton syringe (30). Injections were delivered at a rate of 0.5 µl/minute and the glass needle remained at the injection site for 5 minutes before retraction to avoid backflow. After surgery, rodents were placed in a cage on a heating pad and monitored ambulatory and then administered buprenorphine analgesia (i.p., 0.05 mg/kg) every 8-12 hours for up to 2 days.

### SMM-189 formulation

SMM-189 was provided by Dr. Bob Moore II and formulated with 15% ethanol (v/v), 15% cremophor (w/v) and 70% saline (v/v). In brief, SMM-189 was dissolved in 15% ethanol, cremophor was weighed in a separate vial and dissolved SMM-189 slowly added and mixed gently until combined. Saline was slowly added and mixed until achieving a homogenous solution. The final solution was sterile filtered with a syringe 0.22µm filter and stored at 4°C in amber sterile-glass vials for up to 2 weeks. Vehicle treatments consisted of the same concentrations of ethanol, cremophor and saline. Daily injections were administered by research staff blinded to cohort treatment history.

### Blood and CSF collection

Blood was collected from the tail vein at baseline and at baseline and every 4 weeks post-AAV (∼2 hours post-treatment injection) for plasma to measure SMM-189 levels in low and high cohorts and for PBMCs to immunophenotype circulating immune cells in the high cohort animals. Animals were placed in an induction box and anesthetized with 3% isofluorane and maintained on 1-2% isofluorane for the remainder of the procedure. Tails were cleaned with ethanol and approximately 400uL of blood drawn from the lateral tail vein using a low dead space 25ga needle and syringe and blood immediately placed into an EDTA tube. Body temperature was maintained using a heating pad during blood draw and while recovering from anesthesia.

CSF was collected at endpoint from low cohort animals. In brief, after euthasol injections, rats were placed in a stereotaxic frame and the head fixed downwards. An incision was made to expose the muscle layer over the cisterna magna. A sterile pulled micropipettes attached to tubing and a syringe punctured into the cisterna magna and clean CSF was aspirated and collected in a 0.2mL Eppendorf and stored at −80°C. CSF was not collected from the high cohort rats because brains were extracted quickly and CSF collection may have compromised brain immune cell composition.

### Mass Spectrometry and Assessment of SMM-189 in Plasma and CSF

SMM-189 and *d*^5^-SMM-189 were synthesized as previously described (31). A total of 50 μL aliquots of plasma or CSF were prepared by protein precipitation with 100 μL acetonitrile (spiked with the internal standard *d*^5^-SMM-189) followed by centrifugation at 10,000*g* for 10 min at 4°C. Chromatographic separation of the supernatant was performed using a Zorbax SB-C18 3.5 µm 4.6×150 mm and a Shimadzu HPLC Nexera XR with Solvent Delivery Module LC-20ADXR, CTO-20AC prominence column oven (temperature 40°C), a DGU-20A5R Degassing Unit, and a SIL-20ACXR autosampler. The aqueous mobile phase (A) was 95% water and 5% acetonitrile (B) was 100% acetonitrile. The flow rate was set to 500 µL/min and the gradient elution initial 50% B, 0.5 to 3min 50% to 100%B, 6 to 6.1min 100% to 50% B, 6.5min stop with a column wash using methanol:water (1:1). Sample analysis was performed using SCIEX Triple Quad 5500 tandem mass spectrometer (Framingham, MA) operated in the negative ion mode at ion spray voltage −4500V and temperature 600°C. Multiple reaction monitoring mode using the compound-specific mass transfers of m/z 357 → 185 for SMM-189, and m/z 362 → 185 for *d*^5^-SMM-189 was used for analysis. A calibration curve ranging from 1 to 100 ng/mL was constructed and validated with spiked samples of rat plasma. The peak area ratios of analyte to the internal standard were linear over the tested concentration range.

### Immune cell isolation

PBMCs were isolated from blood of low cohort rats at euthanasia (8 weeks) for qPCR analysis of immune genes and from blood of high cohort rats at baseline, 4 and 8 weeks after AAV2/5-human wild-type Asyn and evaluated with flow cytometry for a longitudinal immunophenotype assessment. Red blood cells were lysed using 1x RBC lysis buffer (Biolegend Cat#420301) for 5 minutes in the dark. Lysis buffer was neutralized by adding 1x HBSS (Gibco Cat#14175095). Cells were pelleted by centrifugation at 400*g* for 5 minutes, supernatant removed, and cells washed with HBSS. For the high cohort, cells were resuspended in 200uL of PBS and transferred to a 96-well v-bottom plate for flow cytometry staining. For low cohort, 10,000 cells were lysed in 600uL RLT lysis buffer (Qiagen Cat#79216) + 1% BME, further homogenized by centrifugation through a QiaShredder column (2 minutes at 21,130*g*). Lysate was then stored at - 80°C and later processed for RNA extraction.

Brains were collected from animals in the high cohort at either 4 or 8 weeks after AAV2/5-human wild-type Asyn and brain immune cells were isolated from individual hemispheres as previously described (32). Hemispheres were finely cut in RPMI 1640 media (Gibco Cat#11875-093) with a scalpel blade and enzymatically dissociated with collagenase VIII (1.4U/mL; Sigma Cat#C2139) and DNAse1 (1mg/mL; Sigma Cat#DN25) in 37°C for 15 minutes. Enzymes were neutralized with 10%FBS (heat inactivated, Atlanta Biologicals Cat#S11150) in RPMI 1640 complete media and centrifuged 400*g* at room temperature. Tissue was resuspended in 1x HBSS and mechanically dissociated using a fire-polished Pasteur pipette until a cell suspension was generated. The suspension was passed through a 70μM cell strainer and strainers rinsed twice with 1x HBSS. To remove myelin, the strained suspension was centrifuged at 300*g* at 4°C for 5 minutes and resuspended in 37% percoll and layered between 70% and 30% percoll gradients. The samples were centrifuged at 400*g* for 30 minutes at room temperature with the brake disabled. The top myelin layer was aspirated and demyelinated immune cells were collected from the 37% and 70% percoll interface with a pipette and washed with 1x HBSS twice, resuspended in PBS and placed in a 96-well v-bottom plate for flow cytometry staining.

### RNA extraction and qPCR

Total RNA was extracted from the frontal cortex and from PBMCs of low cohort animals and standardized across all samples and groups (33). In short, tissue was suspended in Trizol and homogenized with a TissueLyser (Retsch). Homogenate was incubated with chloroform and RNA separated by centrifugation of 113,523*g* for 15 minutes at 4°C. RNA was further isolated with a RNAeasy mini kit (Qiagen). The RNA (1μg) was then reverse transcribed to cDNA and analyzed with primers (Integrated DNA Technologies) for pro-inflammatory and anti-inflammatory immune markers and CB2 (**Table 1**) by qRT-PCR using protocols previously published (33). Resulting threshold cycle (Ct) values were analyzed by the 2-ΔΔCt method using the AAV2/5-hαsyn + vehicle as the control group and normalized to GAPDH for PBMCs and HPRT1 for frontal cortex as the housekeeping genes.

**Table 1.**
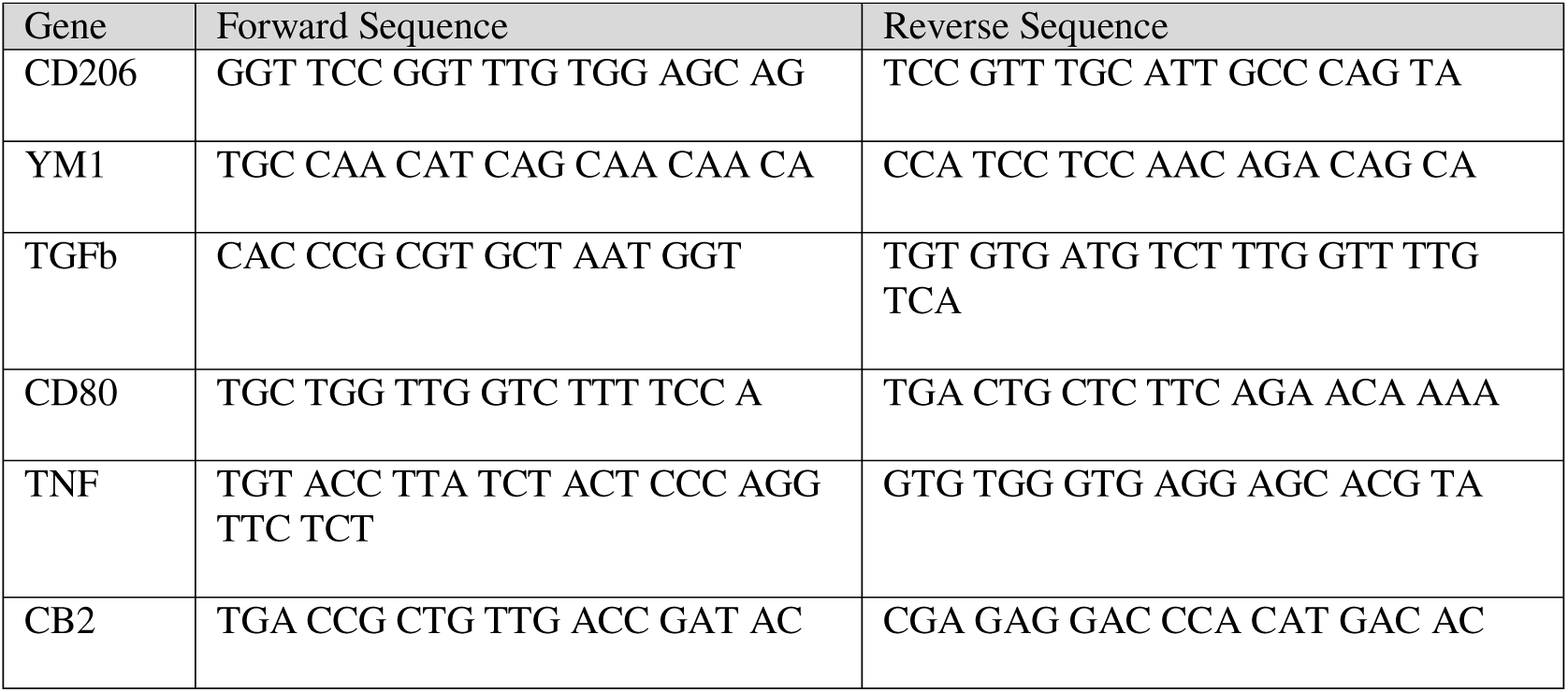
List of rat primers.

### Multicolor flow cytometry

Peripheral and central immune cell staining was performed similarly as previously published protocols (32). In brief, immune cells were washed and resuspended in PBS, then incubated for 30 minutes at room temperature in LIVE/DEAD Fixable Red Dead cell stain (ThermoFisher Scientific Cat#L23102; 1:2000). Following two washes in PBS, Fc receptors were blocked with anti-rat CD32 Fc block diluted 1:100 in FACS buffer (1mM EDTA and 0.1% sodium azide in PBS) for 15 minutes at 4°C. PBMCs were stained for separate myeloid and lymphoid panels and brain immune cells stained with an antibody panel to identify microglia, monocytes and lymphocytes (**Table S1-3**). Fluorophore conjugated antibodies diluted with FACS buffer were incubated at 4°C for 20 minutes. Cells stained for surface myeloid markers were fixed in 1% PFA for 25 minutes and washed in FACS.

PBMCs stained with lymphoid panel were further probed intracellularly for transcriptional marker Foxp3 to label helper Tcells. Here, cells were incubated for 30 minutes in fixation/permeabilization working solution (eBioscience Cat#00-5523-00), washed in 1x permeabilization buffer, additionally blocked with 2% mouse serum for 15 minutes and anti-rat foxp3-AF647 added to block for an additional 30-minute incubation. Then washed and resuspended in FACs buffer. All samples had 10uL of AccuCheck Counting beads (ThermoFisher Cat#PCB100) added to final suspension and cells were run on a LSRII Instrument (BD Biosciences, Franklin Lakes, NJ) with FACSDiva software (BD Biosciences). Results were analyzed using FlowJo 10.8.1 software and brain immune populations of monocytes (classical and non-classical), lymphocytes (CD4+ and CD8+) or microglia and PBMC populations defined by specific strategies (**Figure S1-S3**)(34–36).

For PBMC analysis, raw frequencies were normalized across serial runs against vehicle-treated values using the following equation: Raw frequency / (average vehicle frequencies of specific run / average baseline frequencies). For brain immune cell analysis of 8 weeks, samples could not be collected and ran the same day so frequencies were normalized across treatment group at each timepoint using the following equation: raw frequency / (average treatment frequency of specific run 1 / average treatment frequency of specific run 2). Brain immune cells measurement from 4-week collection were not normalized as flow cytometry was performed on all samples at the same time.

### Immunohistochemistry

Brains were post-fixed in 4%PFA for 48 hours and transferred to 30% sucrose and cut at 40μm on a freezing microtome. Immunohistochemical stains were performed as previously published with antibodies listed in **Table S4** (37). In brief, endogenous peroxidase was quenched with 3% hydrogen peroxide, blocked with 8% normal serum, avidin (Vector Cat#SP-2001) and 0.1% Triton-X and incubated overnight at 4°C with biotin (Vectorlabs Cat#SP-2001), normal serum and primary antibodies specific for Asyn (Clone 4B12; Biolegend Cat#807801), pSer129 Asyn (Clone EP1536Y; Abcam Cat#AB51253), CD68 (Serotec Cat#MCA341R), MHCII (Biorad Cat#MCA46GA), and CD163 (Biorad Cat#MCA342A). Tissue was incubated in appropriate biotinylated secondary antibodies (1:200), amplified with Vectastain ABC kit (Vectorlabs Cat#PK-6100) and developed with diaminobenzidine tablets or diaminobenzidine drop kit with nickel (Vector Cat#SK-4100) for CD68 immunostaining.

Proteinase-K resistant Asyn staining was completed with one piece of matched nigral tissue mounted and dried onto positive charged slides. Once rehydrated, a hydrophobic barrier was drawn around the tissue and proteinase K (1:1000 in PBS) incubated for 30 minutes at room temperature. Proteinase k was cleared with PBS and the remaining staining methods were identical to those listed above with antibodies including Asyn Clone 4B12 1:5000 and horse anti-mouse biotinylated secondary 1:500.

### Immunofluorescence

Tissue was blocked with 5% normal serum in TBS + 0.25% Triton-X and further incubated overnight at 4°C with primary antibodies specific for IBA1 (Wako Cat#019-19741) and tyrosine hydroxylase (Immunostar Cat#22941) to identify microglia in the substantia nigra or tyrosine hydroxylase (Millipore AB152) and Asyn (Clone 4B12; Biolegend Cat#807801) to evaluate accuracy of stereotaxic injection targets. Tissue was incubated in secondary antibodies conjugated to appropriate fluorophores for 2 hours at room temperature and mounted tissue with Vectashield, an aqueous mounting media with DAPI (Vectorlabs Cat#H-2000).

### Histological analysis and quantification

Low cohort animals were assessed by an experimenter blinded to the treatment history for rigorous and proper assessment of unilateral targeting of AAV2/5-human wild-type Asyn into the SN using immunofluorescent analysis of human Asyn and tyrosine hydroxylase (**Figure S4**). Low cohort animals were excluded from analysis when poor stereotaxic targeting occurred, including missing human Asyn expression throughout the rostral-caudal axis of the SN or if expression was seen only in lateral SN. A total of 6 animals in the low cohorts were removed and the final numbers are reflective in the animals section of the Methods.

IBA1-positive total immune cell counts and area of immunoreactivity were quantified using Nikon NIS elements software. Four fluorescent z-stacks of IBA1 stained nigra were collected from images of lesioned and non-lesioned nigra from targeted low cohort animals using a Nikon Eclipse 90i microscope and maximum project images were used for quantification. CD68-ir and CD163-ir was quantified from two to three microphotographs from each animal throughout the substantia nigra using ImageJ particle count analysis (38). Microphotographs of the nigra immunochemically stained for either Asyn, pSer129 Asyn and proteinase K-resistant Asyn were quantified using Image J (version 1.54f). Regions of Interest (ROIs) were drawn on two to three images throughout the nigra and optical density measurements collected and averaged for each animal. Measurements were normalized to the vehicle group so that data from all cohorts could be combined.

### SDS-PAGE and immunoblotting

Striatal tissue from successfully targeted cohort 1 and cohort 2 rats was homogenized in ice-cold TriZOL lysed using the TissueLyser system. Homogenates were incubated with 20% chloroform and centrifuged at 13,523*g* for 15minutes at 4°C. Protein pellets were washed in methanol twice and resuspended in 1% SDS and heated to 50°C. Total protein concentrations were measured using Pierce BCA Protein Assay Kit and equal amount of protein (10-20μg) mixed with Laemmli sample buffer were loaded in 4-20% TGX precast gels (Biorad) for gel electrophoresis. Protein was transferred to a PVDF membrane (Biorad) using a Trans Blot Turbo Transfer System, the immunoblots were immediately fixed with 0.4% paraformaldehyde (PFA) for 30 minutes and stained for total protein using Revert 700 (Licor) for reliable normalization. Revert staining was captured and measured on the Licor Odyssey at 700nm for 30 seconds.

Non-specific protein binding was blocked with 5% milk and 0.1% Tween-20 in TBS for 1 hour at room temperature, and membranes incubated with primary antibodies overnight at 4°C. The blots were washed with 0.1%TBS-Tween-20, incubated with appropriate horseradish peroxidase-conjugated secondary antibodies (1:2000) at room temperature for 1 hour and developed with SuperSignal West Femto Enhancer or Pico Peroxide solutions under a chemiluminescent detection imaging system (Licor Odyssey). The density of the band was quantified using StudioLite software. Relative expression levels were normalized to its own total protein and to vehicle average expression level.

### In vitro cell culture

The RAW264.7 macrophage cells (ATCC Cat#TIB-71) were grown in sterile media which consisted of DMEM high glucose, 10% FBS, 2mM L-glutamine, 1x Pen-strep and 1x sodium pyruvate in a humidified 5% CO2 incubator at 37°C. RAW264.7 were maintained in 75cm^3^ tissue culture flasks and plated at a density of 1×10^4^ cells in triplicate in 96-well plates for assays. Cells were stimulated with either lipopolysaccharide (LPS O111:B4; Sigma Cat#L2630) at 100ng/mL or 1000ng/mL (750EU/mL or 7500 EU/mL, respectively). One hour prior to this stimulation the cells were pre-treated with the CB2 inverse agonist SMM-189 at 3uM or 7uM or its vehicle. Specifically for cell culture experiments, SMM-189 was formulated with 40% ethanol (v/v), 20% tween-20 (v/v) and 40% saline (v/v). A phagocytosis assay was conducted 24 hours after LPS.

### Phagocytosis assay

The phagocytosis assay was performed using pHrodo green *E. coli* bioparticles (Invitrogen Cat# P35366) according to the manufacturer’s instructions. Five images per well were then collected every 30 minutes for 24 hours on an EVOS M7000 microscope (Invitrogen, ThermoFisher Scientific). The cells were maintained for this duration using an on-stage incubator kept at 37°C and 5% CO_2_. Analysis of the collected images was performed using EVOS Celleste software (version 5.0), which counted the cells and then measured the integrated optical density of fluorescence within each well over the 24-hour time course. Analysis was conducted on the average integrated optical density across triplicate wells. Results were consistent across duplicate experimental replicates.

### Statistics

Statistical analysis was conducted using GraphPad Prism software (version9.5.1). *In vivo* histological, western and transcriptional analyses were analyzed using parametric unpaired t-tests to determine the statistical difference between SMM-189 and vehicle-treated groups. Flow cytometry data of brain immune cells was analyzed using a two-way ANOVA (factors of hemisphere and treatment) and multiple comparisons performed using uncorrected Fishers LSD, while PBMC serial flow data was analyzed using a fixed-effect model (factors of time and treatment) and multiple comparisons performed using uncorrected Fishers LSD. Cell experiments were analyzed using a one-way ANOVA to determine differences between treatment groups across the entire timed experiment using a Dunnet’s multiple comparison test.

## Results

### Phosphorylated Asyn in substantia nigra is decreased by modulation of CB2 with SMM-189

To assess the effect of SMM-189 on Asyn aggregation, midbrains from the low cohorts were immunohistochemically stained for human phosphorylated (pS129) Asyn (clone EP1536Y; abcam Cat#ab512253). This phosphorylation site is pathologically relevant as it has been found in Lewy bodies in postmortem samples (39, 40) and has been shown to accelerate aggregation of Asyn in vitro and in vivo (41, 42), suggesting that pS129 is linked to aggregation. Optical density analysis of the substantia nigra revealed reduced pS129-immunoreactivity in SMM-189-treated animals when normalized against vehicle-treated animals (t(23)=3.353, p=0.0023; **Figure 2A,B**). Total and proteinase-K resistant Asyn was also evaluated in the substantia nigra and found that while levels of total human Asyn (Biolegend, clone 4B12) were not different between treatments, SMM-189 decreased PK resistant Asyn compared to vehicle (t(10)=2.303, p=0.044; **Figure 2C-F**) when normalized to vehicle nigra measurements. Consistent with histological analyses of pS129 and total Asyn in the nigra, western analysis of the striatum ipsilateral to the overexpression of human Asyn demonstrated reduced pS129 levels in SMM-189-treated rats (t(15)=3.710, p=0.0021), and no differences in total human Asyn levels were found between treatments (p=0.2133; **Figure 2G,H**).

**Figure 2.**
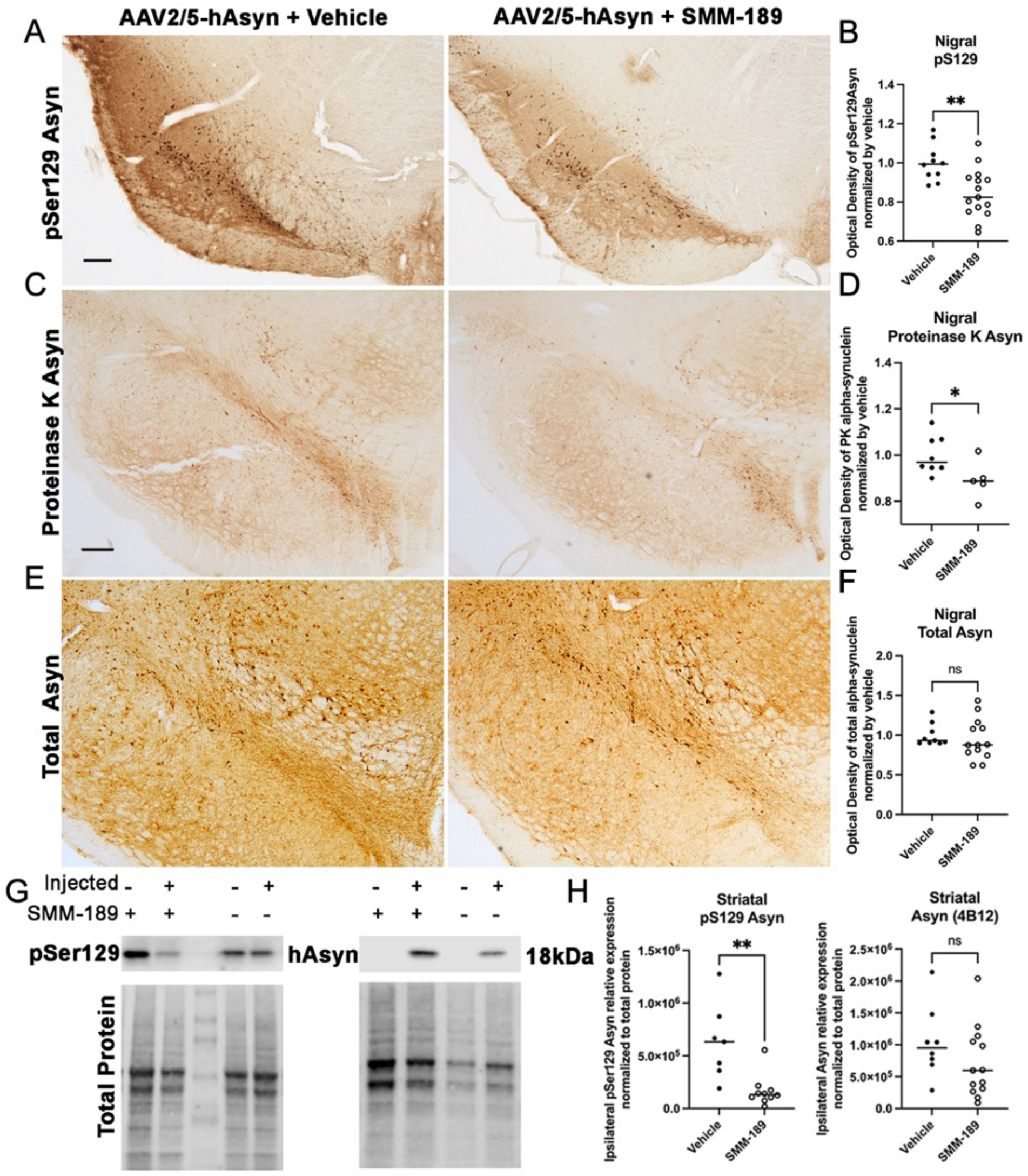
Modulation of CB2 with SMM-189 decreases accumulation of pSer129 alpha-synuclein in the lesioned nigra. Representative images of pSer129 Asyn (A, scale bar=250μm), proteinase K-insoluble Asyn (C), and total human Asyn (E) in rats that received AAV2/5-hAsyn and treated with either vehicle or SMM-189. Quantification of low cohort rats treated with SMM-189 normalized to vehicle showed reduced pSer129 Asyn (abcam ab512253, B) compared to vehicle and reduced intensity of proteinase K-insoluble Asyn (D). Total human Asyn did not differ between groups when evaluating the optical density of Asyn-ir within the lesioned nigra normalized by vehicle (F). Representative western immunoblots of pSer129 and total human synuclein from striatal lysates and their corresponding total protein used for normalization (G). Quantification of western bands of hemispheres ipsilateral to the AAV2/5-hAsyn injections demonstrate that pSer129 was significantly reduced following SMM-189 treatment, but human Asyn was not different between treatments (H). *p<0.05, **p<0.01

### SMM-189 measurements in CSF confirm penetration into rat brain

Next, we evaluated the weights and levels of SMM-189 from plasma of rats from low and high cohorts. Rats weight increased overtime (p<0.0001) as expected in both low and high cohorts with no significant differences between treatment groups (**Figure S5A,D**). Significant increases were detected in the level of plasma SMM-189 in animals receiving SMM-189 treatment in low cohort (treatment F(1,7)=16.46, p=0.0048; time F(2,14) = 8.647, p=0.0036) at 4 (p=0.0222) and 8 weeks (p=0.0001) compared to vehicle-treated rats (**Figure S5B**). Levels of SMM-189 remained low in the CSF with an average of 0.44±0.22SEM ng/mL (**Figure S5E**) and a brain to plasma ratio average of 0.032±0.02SEM. This is in contrast to the brain to plasma ratios measured for SMM-189 in murine PK studies wherein the brain to plasma ratio of SMM-189 was 0.74 following i.p. injection and 1.63 after p.o. administration (**Table 2**). Low levels of drug in the CSF are presumed to reflect the extensive portioning of SMM-189 (clogP value of ∼5.26) into the brain tissue based on high penetration measured in murine PK studies. Similar to low cohort, SMM-189 plasma levels in the high cohort increased with time (treatment: F(1,68), p<0.0001; time F(2,68)=22.24, p<0.0001) in SMM-189-treated rats compared to vehicle at 4 (p<0.0001) and 8 weeks (p<0.0001; **Figure S5E**). CSF was not collected from high cohort rats.

**Table 2.** Pharmacokinetic results of SMM-189 peripheral administration in mice.

| Matrix | Route | Dose (mg/kg) | Tmax (hr) | Cmax (ng/mL) | Brain/Plasma Ratio (Cmax) |
| --- | --- | --- | --- | --- | --- |
| Plasma | i.p. | 6 | 0.50 | 277.07 |  |
|  | p.o. | 6 | 0.25 | 43.15 |  |
| Brain | i.p. | 6 | 0.50 | 204.64 | 0.74 |
|  | p.o. | 6 | 0.25 | 70.34 | 1.63 |

### SMM-189 promotes an anti-inflammatory environment and shifts immune cell gene expression toward a wound healing phenotype in the CNS

To evaluate the effects of CB2 modulation on neuroinflammation, nigral sections were immunohistochemically examined for immune cell markers. SMM-189 did not change microglial IBA1+ numbers or area (**Figure 3A,B**). IBA1 analysis demonstrated lesion effects with significantly increased counts (F(1,22)=18.72, p=0.003) and area (F(1,22)=22.94, p<0.0001) in the Asyn-lesioned nigra compared to the uninjected side. Notably, the lesion effect on IBA1 was lessened in SMM-189-treated rats compared to vehicle-treated rats, potentially highlighting the ability for SMM-189 to dampen Asyn-activated IBA1+ brain myeloid cells. In the striatum, western blot analysis revealed no differences in myeloid marker IBA1 and astrocyte marker GFAP expression levels across treatments (**Figure S6**). SMM-189 did promote an anti-inflammatory environment with increased nigral CD68-immunoreactive particle counts (t(15)=3.413, p=0.0039) in the substantia nigra injected with AAV-wt-hAsyn compared to vehicle treated conditions (**Figure 3C,D**). CD68 is a lysosomal marker and often a surrogate marker for phagocytic innate immune cells, suggesting an increase in phagocytic wound-healing cells in SMM-189-treated brains. CD163, an innate-immune marker identifying cells from the monocyte/macrophage lineage, was also histologically evaluated in the nigra and found to be decreased in SMM-189-treated rats compared to vehicle-treated animals (t(7)=5.172, p=0.0013).

**Figure 3.**
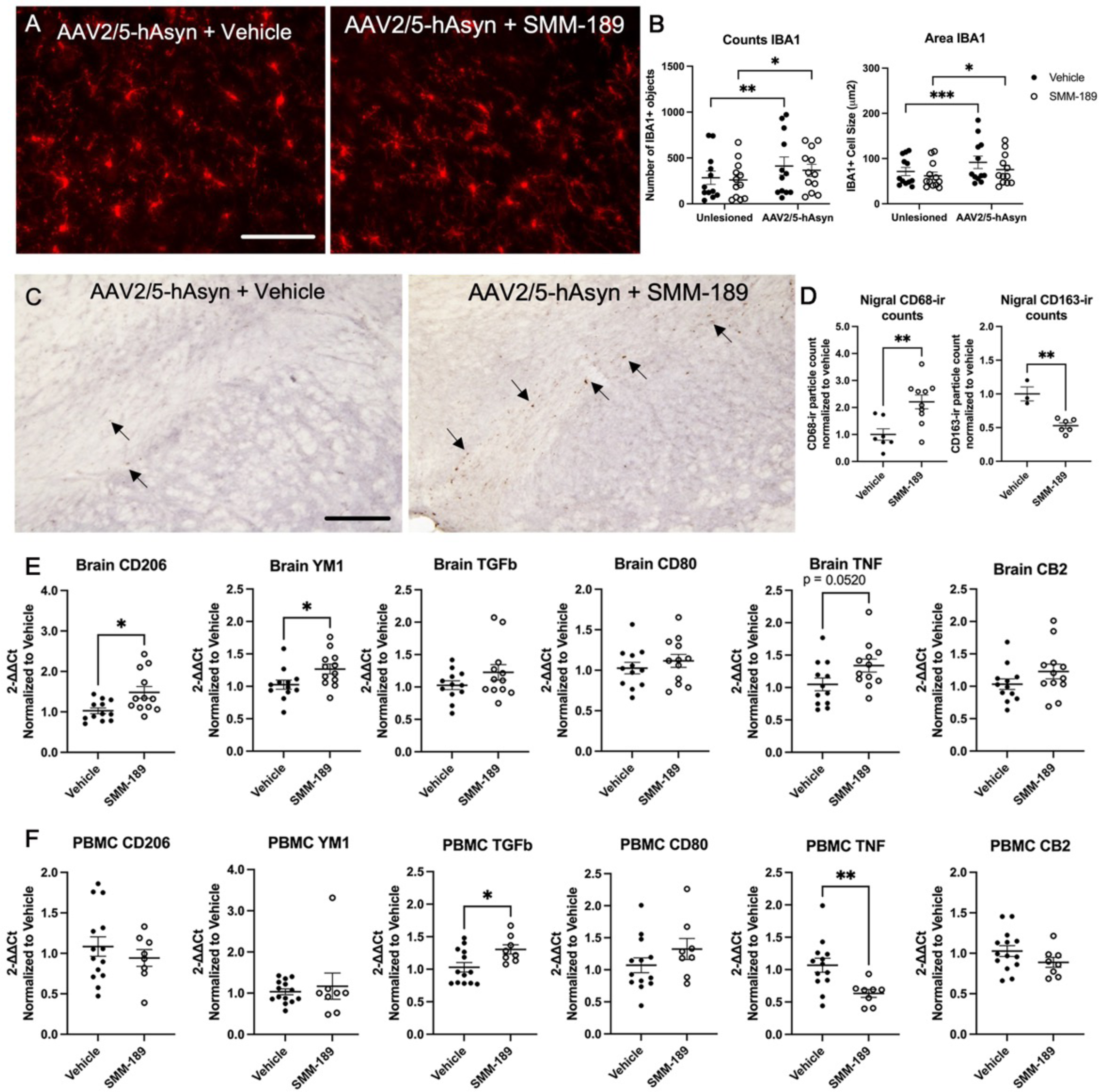
Modulation of CB2 with SMM-189 modifies central neuroinflammation. Representative microphotographs of midline midbrain sections immunostained for IBA1 (A). Quantification of IBA1+ immunofluorescent staining reveals significant lesion effect based on count and area for each treatment group. Scale bar = 100μm (B). Representative images of midbrain sections immunostained for CD68, scale bar = 500μm (C). Arrows highlight the CD68+ cells included in the quantification. CD68 quantification using ImageJ particle counting and normalized to vehicle found significant increases in SMM-189 compared to vehicle-treated rats, while CD163 quantification within the lesioned nigra was decreased by SMM-189 treatment (D). The mRNA expression of inflammatory cytokines CD206, YM1, TGFβ, CD80, TNF, CB2 are displayed for both frontal cortex or brain (E) and PBMCs (F). Relative PBMC mRNA abundance was normalized to GAPDH, frontal cortex to HPRT1. *p<0.05, **p<0.01, ***p<0.001

To further assess inflammation, cytokines transcript levels were evaluated in the frontal cortex brain tissue and PBMCs of low cohort rats by qRTPCR. SMM-189-treated rats displayed increased M2 alternative activation markers YM1 and CD206 gene expression in the frontal cortex and TGFβ in PBMCs relative to vehicle-treated animals (**Figure 3E,F**). Endpoint PBMCs also demonstrated decreased pro-inflammatory cytokine TNF transcript after SMM-189 treatment. CB2 gene expression itself was not different between groups confirming that the target is present in immune cells and brain.

### SMM-189 induced significant time-dependent alterations in profiles of brain-infiltrating immune cells and in peripheral monocyte populations

Next, to investigate the effects of SMM-189 on the infiltration of immune cells from the periphery, brain immune cells were isolated from the high cohort by enzymatic digestion and percoll gradient separation at both 4- and 8-weeks post-nigral injections. Flow cytometry analysis at 4 weeks (**Figure 4**) revealed that SMM-189 treatment globally reduced CD172a+CD11b+ monocyte frequency (treatment: F(1,18)=9.310, p=0.0069) with no significant changes detected in more defined classical or non-classical monocyte subset frequencies. However, the mean fluorescent intensity (MFI) of the signal regulatory protein A (SIRPa) CD172a, an inhibitory receptor that binds to “do not eat me” ligand CD47), was dependent on the AAV injection on CD43-classical monocytes (interaction: F(1,18)=5.80, p=0.027), such that the unlesioned hemisphere displayed elevated CD172a in vehicle-treated rats compared to SMM-189 and compared to its own vehicle-treated lesioned hemisphere. No microglia or CD3+ lymphocytes population differences were identified at 4 weeks, however the MFI of CD11b (activation marker) microglia was reduced in unlesioned hemispheres of SMM-189-treated rats compared to control (p=0.0151) and compared to its own lesioned hemisphere (p=0.0124). Evaluation of T cell subsets at 4 weeks identified an overall reduction in the frequencies of CD4+ T cells in brains of SMM-189-treated rats (treatment: F(1,18)=5.46, p=0.0311). Although CD8 T cell frequencies were not significantly different, the CD4 to CD8 ratio was found to be decreased at 4 weeks in the hemispheres administered AAV2/5-human wild-type Asyn injections when exposed to SMM-189 (p=0.0443).

**Figure 4.**
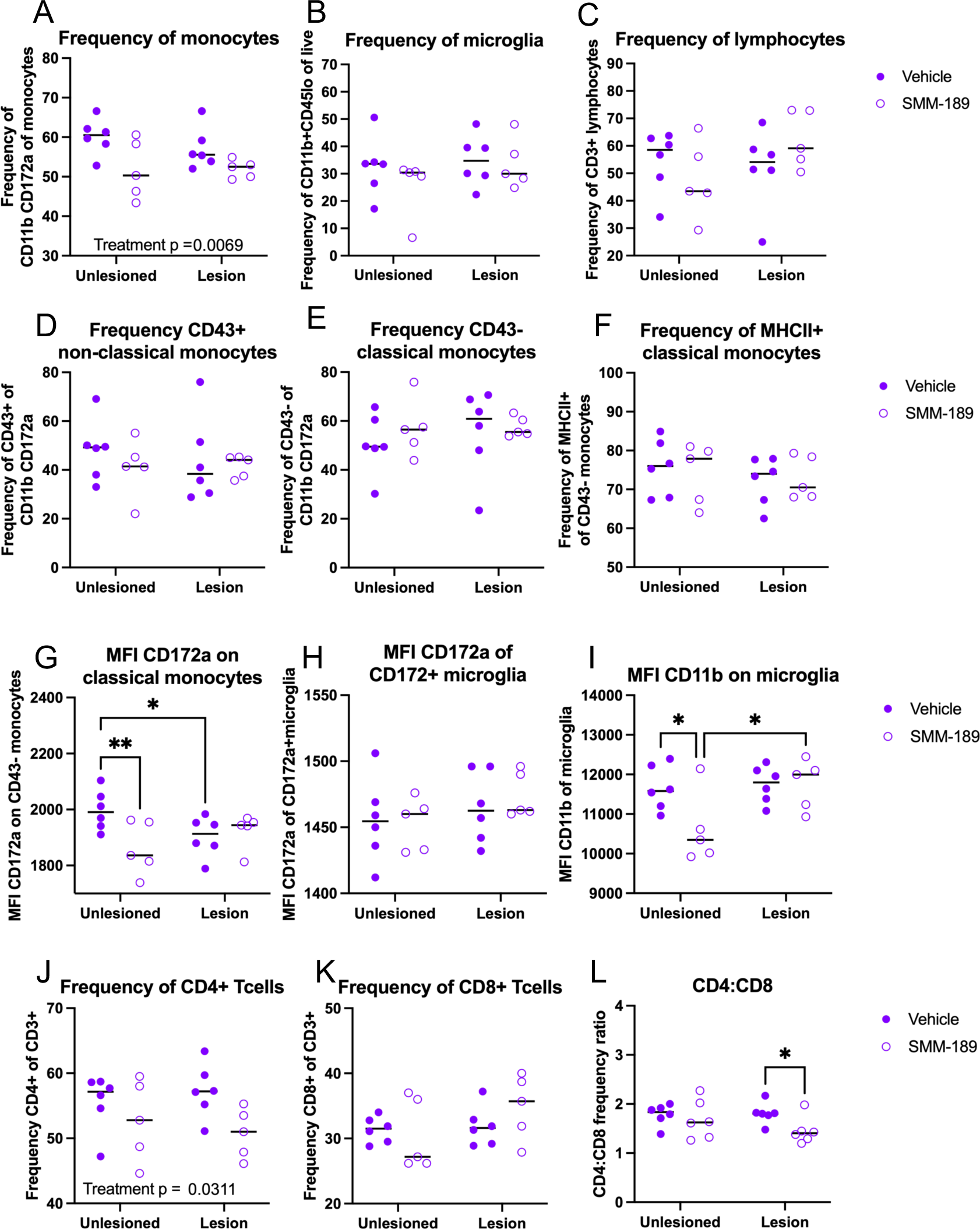
Modulation of CB2 with SMM-189 globally reduces infiltrating monocytes and brain myeloid cell activation at 4 weeks. Brain immune cells from high cohort rats were isolated from lesioned (+human Asyn) and unlesioned (-human Asyn) rat hemispheres at 4 weeks and monocyte, lymphocyte and microglia populations initially identified from CD45 and CD11b expression and further gated into more distinct populations. Quantification of immune cell population frequencies are shown here for monocytes (CD11b+CD172a+), microglia (CD45loCD11b+) and lymphocytes (CD3+) of their parent gate (A-C), with significant reductions in monocyte frequencies in SMM-189 rats (A) and no differences in microglia (B) or CD3+ lymphocytes (C). Quantification of non-classical (CD43+CD172a+), classical (CD43-CD172+) monocytes and MCHII+ classical monocytes show no differences across groups (D-F). Mean fluorescent intensity (MFI) was evaluated for monocyte and microglia populations and found alterd CD172a on classical monocytes (G) with no effect of CD172a on microglia (H). MFI of CD11b was reduced in unlesioned hemispheres of those with SMM-189 treatment compared to vehicle and also found significantly lower than SMM-189 lesioned hemispheres (I). T cell subsets were quantified and found that CD4+ T cells were reduced in lesioned hemispheres of SMM-189 rats compared to vehicle (J). Although no differences were seen in CD8+ T cells (K), a reduced CD4:CD8 ratio was identified in the presence of Asyn (L). *p<0.05, **p<0.01

Interestingly, 8 weeks after AAV injections, SMM-189 shifted brain immune cells frequencies in the opposite manner than at 4 weeks (**Figure 5**). The monocyte frequency (CD172a+CD11b+) was elevated in brains of SMM-189-treated rats (treatment: F(1,24)=11.71, p=0.0022) where multiple comparisons find elevations in both the lesioned hemisphere (p=0.0145) and the unlesioned hemisphere (p=0.0372) from SMM-189 treatment compared to vehicle. Although more defined monocyte populations (classical and non-classical) were not found different between treatments, the intensity of MHCII on classical monocytes was increased in the lesioned hemisphere in SMM-189-treated rats compared to vehicle (p=0.0172). Similar to parent populations at 4 weeks, no differences were found in microglia or CD3+ lymphocytes across groups. However, SMM-189 globally elevated microglial MFI of CD172a (treatment: F(1,24)=5.659, p=0.0257) and activation maker CD11b (treatment: F(1,24)=7.611, p=0.0109) with significant elevation in the non-lesioned hemisphere from multiple comparisons (p=0.0162). Furthermore, independent of the presence of Asyn, CD4+ lymphocytes globally increased (treatment: F(1,24)=17.52, p=0.0003), while CD8+ lymphocytes decreased (treatment: F(1,24)=23.79, p<0.0001) in SMM-189-treated animal brains compared to vehicle, resulting in increased CD4 to CD8 ratio in the brains of SMM-189-treated rats 8 weeks after AAV2/5-human wild-type Asyn injections.

**Figure 5.**
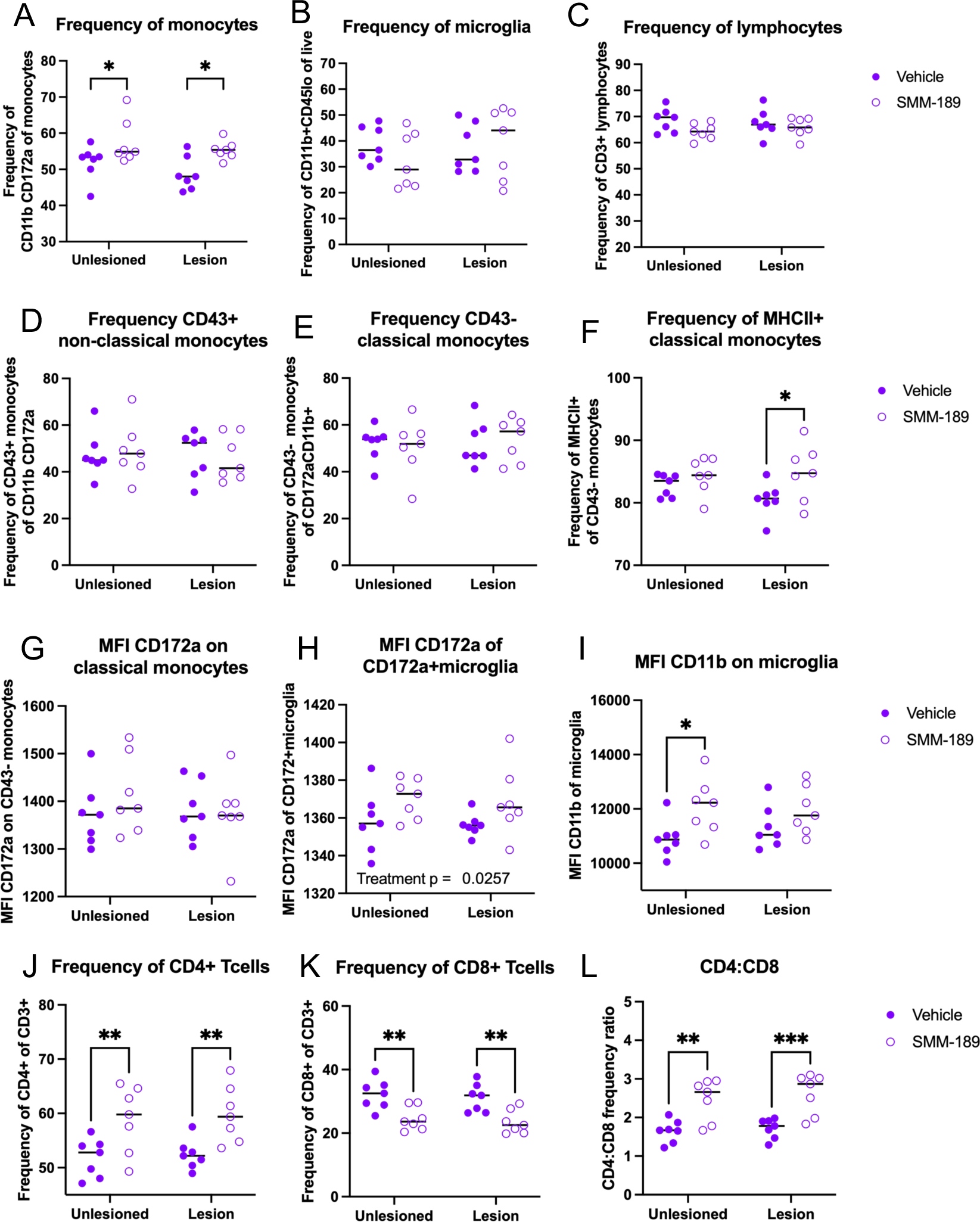
Modulation of CB2 with SMM-189 promotes monocyte infiltration and increased innate and adaptive immune cell activation in the brain. Brain immune cells from high cohort rats were isolated from lesioned-(+human Asyn) and unlesioned (-human Asyn) hemispheres at 8 weeks and monocyte, lymphocyte and microglia populations initially identified using CD45 and CD11b expression and further gated into more distinct populations. Quantification of immune cell population frequencies are shown here for microglia (CD45loCD11b+), monocytes (CD11b+CD172a+) and lymphocytes (CD3+) of their parent gate (A-C). Monocyte frequencies were increased in the brains of SMM-189-treated animals compared to vehicle-treated animals in both hemispheres (A), but no differences in microglia (B) or CD3+ lymphocytes (C). Quantification of non-classical (CD43+CD172a+), classical (CD43-CD172+) monocytes show no differences across groups (D-E), yet treatment with SMM-189 increased the frequency of MCHII+ classical monocytes in the presence of Asyn compared to treatment with vehicle (F). Mean fluorescent intensity (MFI) of CD172a on classical monocytes was not different between conditions (G), but was globally increased on CD172a+microglia from SMM-189-treated rats (H). MFI of CD11b on microglia was increased in unlesioned hemispheres of those with SMM-189 treatment compared to vehicle (I). CD4 and CD8 T cell subsets were quantified and found that CD4+ T cells were increased (J), while CD8+ T cells were reduced (K) in both hemispheres of SMM-189-treated rats compared to vehicle, resulting in increased CD4:CD8 ratios across treatment groups (L). *p<0.05, **p<0.01, ***p<0.001

In the periphery, SMM-189 reduced circulating classically-activated monocytes (CD43-His48+) as determined by flow cytometry (treatment: F(1,24)=5.093, p=0.0334) with a multiple comparisons demonstrating a significant decrease at weeks 4 (p=0.0465) and 8 (p=0.0384) in PBMCs of SMM-189-treated rats compared to vehicle-treated animals (**Figure 6**). Yet the amount of MHCII as determined by MFI was increased on CD43-His48+ classical monocytes at 4 weeks after SMM-189 treatment compared to control (p=0.0328). Furthermore, at 8 weeks increases in the wound-healing non-classical monocytes (CD43+; p=0.0466), which circulate and patrol in search of injury, and decreases in granulocytes (p=0.0355) was found in SMM-189 compared to vehicle-treated using multiple comparisons analysis. Lymphocyte frequencies of PBMCs did not reveal significant differences in CD4 or CD8 populations (or the CD4 to CD8 ratio) between treatments, however, animals treated with SMM-189 had elevated levels of Tregs (CD4+Foxp3+, p=0.004; and further defined as CD4+Foxp3+CD25+, p=0.0145) at 8 weeks compared to their own baseline frequencies (**Figure S3**). Immune cell populations were also defined in deep cervical lymph nodes (DCLN) and splenocytes of high cohort rats at the 8-week endpoint to evaluate the influence of peripheral SMM-189 treatments on peripheral lymphoid tissues (**Figure S7**). A trend was found in the CD4 to CD8 lymphocyte ratio to be decreased in DCLN (p=0.0503) and spleen (p=0.0962) in SMM-189-treated rats, representing an inverse immune cell composition compared to the brain at 8 weeks. SMM-189 also prompted a trend to increase CD11b+ myeloid splenocytes (p=0.0541) with significant elevation of their MHCII intensity (p=0.0185).

**Figure 6.**
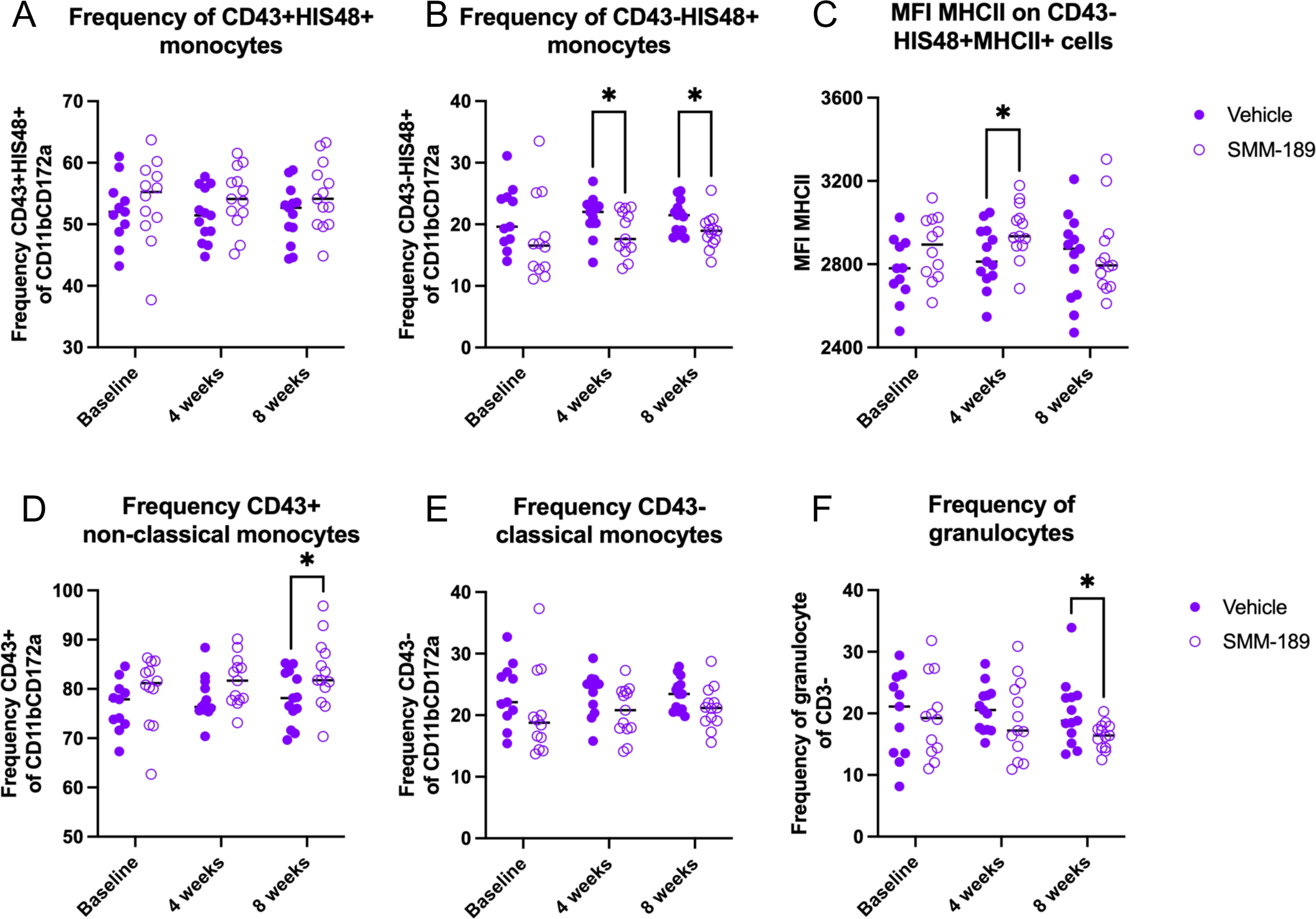
Modulation of CB2 with SMM-189 skews PBMC phenotype and increases frequency of circulating wound-healing non-classical monocytes. Peripheral blood mononuclear cells were isolated from whole blood at baseline, 4 and 8 weeks after AAV2/5-hAsyn injections from rats in the high cohort. SMM-189 treatment resulted in no change in frequency of His48+ non-classical monocytes (A), but decreased circulating His48+ classical monocytes (CD43+CD11b+CD172a+) at 4 and 8 weeks compared to vehicle treatment (B). Mean fluorescent intensity (MFI) of MHCII on His48+ classical monocytes was elevated on PBMCs from SMM-189-treated rats compared to vehicle at 4 weeks (C). Parent gate CD43+ non-classical monocytes demonstrated increased frequency of CD43+ non-classical monocytes (D), but no changes in parent gate CD43-classical monocytes (E). Frequency of granulocytes as defined by CD3- and granularity (SSC) were decreased at 8 weeks from SMM-189 treatment compared to vehicle (F). *p<0.05

### SMM-189 pretreatment enhances LPS-induced phagocytosis in RAW264.7 macrophages

To better understand the effects of SMM-189 on functional uptake and degradation by innate immune cells, a phagocytosis assay was performed using *E. coli* phHrodo bioparticles in RAW264.7 macrophage cells. SMM-189 potentiated the LPS-stimulated response as measured by the increased optical density of GFP expressed in the RAW cells (**Figure 7**). Specifically, when stimulated with 100ng/mL LPS there was a dose dependent response to SMM-189 where pre-treatment with drug at either 3uM or 7uM significantly increased the integrated density of GFP when compared to pre-treatment with vehicle (p=0.0024, p<0.0001, respectively; **Figure 7A**). However, when stimulated with a higher concentration of LPS (1 ug/mL), both 3uM and 7uM doses of SMM-189 intensified phagocytosis of *E. coli* bioparticles compared to vehicle (p<0.0001 and p<0.0001, respectively; **Figure 7B**), suggesting a ceiling effect when cells are highly activated. Regardless of LPS dose stimulation, as a control, GFP expression was significantly restricted with the additional incubation of cytochalasin D, an actin polymerization inhibitor.

**Figure 7.**
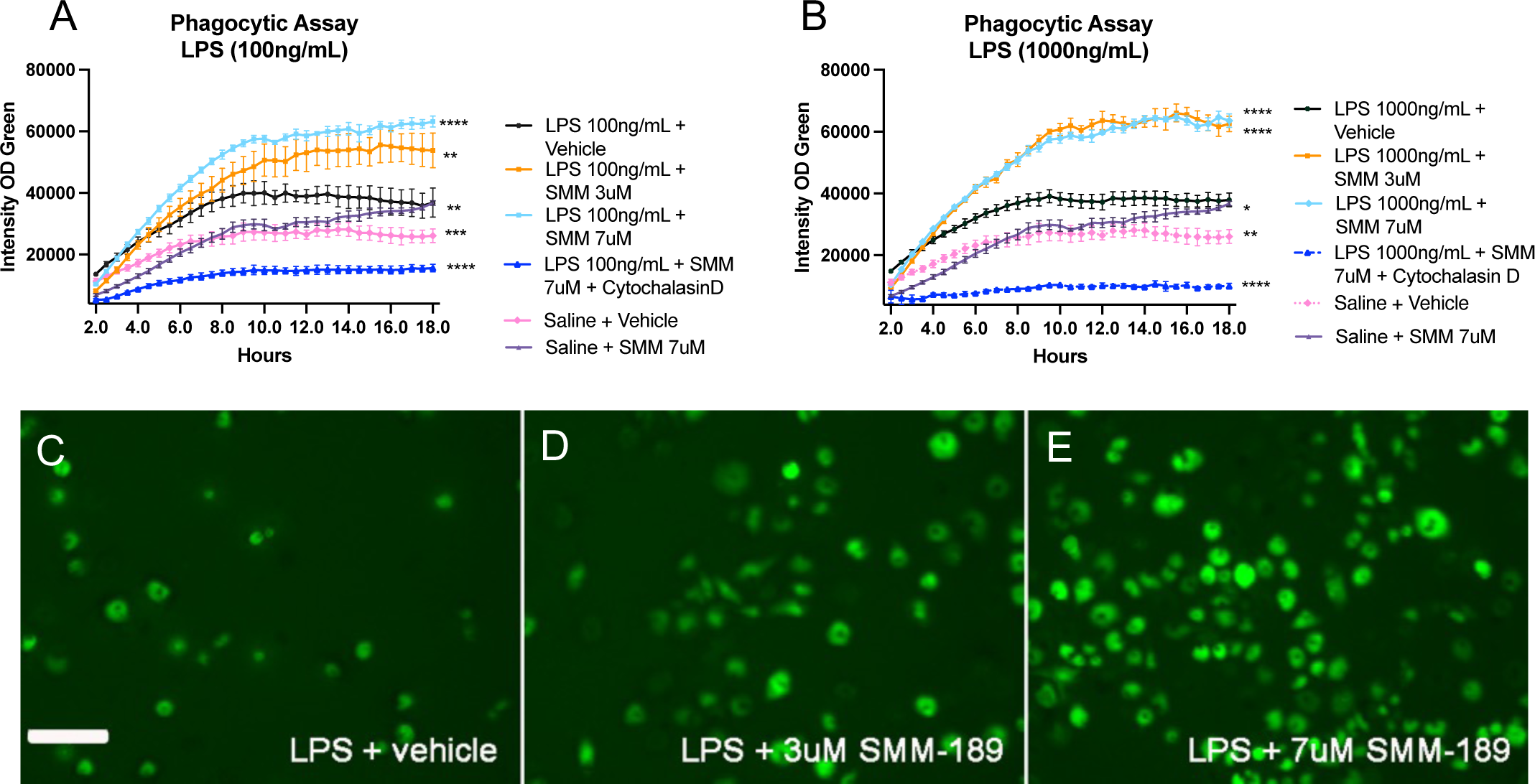
Modulation of CB2 with SMM-189 increases RAW264.7 macrophage phagocytosis after an inflammagen challenge. Quantification of GFP fluorescent intensity in RAW264.7 macrophage cultures challenged with either 100ng/mL (A) or 1000 ng/mL (B) lipopolysaccharide and pre-treated with SMM-189 or vehicle and 24 hours later incubated with pHrodo green E. coli. Treatments are analyzed across time using a one-way ANOVA with Dunnett’s multiple comparisons test. Data are representative of 2 separate experiments. Representative GFP images of fluorescent pHrodo green E. coli at 12 hours for 100ng/mL LPS + vehicle (C), 100ng/mL LPS + 3uM SMM-189 (D), 100ng/mL LPS + 7uM SMM-189 (E) conditions. *p<0.05, **p<0.01, ***p<0.001, ****p<0.0001

## Discussion

Evidence of chronic neuroinflammation and dysfunction of the innate immune system have been described in PD, which has given rise to a growing landscape of therapeutic interventions that target microglia and the peripheral immune system to delay or prevent disease progression (43). The cannabinoid system has become a recent therapeutic target of interest due to its influence on the immune system and the increased legalization across continents of cannabis that directly acts on the cannabinoid receptors. Given the untoward side effects of cannabis intake on psychomotor performance, focus on the immunomodulatory CB2 signaling arm of the pathway represents a significantly novel and more desirable therapeutic direction for the field. This study is the first to evaluate the effects of a CB2-selective ligand on synucleiopathies. The results of the present study demonstrate that CB2 inverse agonism reduces Asyn phosphorylation and aggregation and shifts central and peripheral immune cells toward a wound-healing, phagocytic phenotype in an Asyn rat model of nigral neuroinflammation. These novel results point to a critical role of CB2 in mediating the function of brain immune cells that could be harnessed to prevent or treat synucleinopathies.

Our data demonstrate that SMM-189 treatment reduced phosphorylated Asyn and proteinase-k resistant Asyn, which is in line with other published studies demonstrating that CB2-deficient mouse models resulted in reduced protein aggregation. Specifically, transgenic CB2 knockout mice had significantly decreased Amyloid-β-enriched plaque density in the hippocampus, as measured with Methoxy-XO4 staining, and improved cognitive and learning deficits in mouse models of Alzheimer’s disease-like pathology (44, 45). An additional study found reduced phosphorylated tau levels in insoluble fractions from the hippocampus of CB2-/- mice overexpressing human P301L mutant tau compared to wild-type mice (46). These results would suggest that CB2 plays a role in the removal of aggregated proteins. Although the accumulation of Amyloid-β versus Asyn or tau are differentially localized, extracellular versus intracellular, respectively, evidence from others has supported the extracellular propagation of Asyn from cell-to-cell (47, 48). The reduced Asyn phosphorylation and proteinase-k resistant aggregates found in this study may suggest that modulation of CB2 on immune cells promoted clearance of extracellular Asyn. In the context of aggregated Asyn, only one other study has evaluated the role of CB2 in Asyn clearance and found opposing effects on Asyn accumulation. Feng and colleagues evaluated transgenic CB2-deficient and CB2 wild-type mice treated with human Asyn pre-formed fibrils (PFF) in the nucleus accumbens and found increased area coverage of Asyn 2 hours post-PFF injections in the absence of CB2 (21). Although the study suggests impaired Asyn clearance, authors also report increased intensity of CD68 immunostaining and morphologically more ameboid (less branches) shaped IBA1+ microglia in CB2-/- compared to wild-type mice at the same timepoint, both of which would suggest phagocytic microglial phenotypes. Feng and colleagues did not evaluate insoluble or phosphorylated species of Asyn. Multiple studies have identified phosphorylated Asyn to be critical for Asyn inclusion formation (42, 49), suggesting that the reduced pSer129 in our study may have overall therapeutic value. It is difficult to directly compare our Asyn results to this study because of the many differences in experimental design. Important differences of note include pharmacological inverse agonism in our study compared to transgenic CB2 deficiency in the Feng study, the mode of Asyn overexpression (virus-mediated versus PFF) and the inoculated brain region (substantia nigra versus nucleus accumbens). Microglia are heterogenous across brain regions such that midbrain compared to striatal and pre-frontal cortex microglia have distinct transcriptional profiles enriched for immune-related pathways, which possibly account for opposing results on Asyn (PFF) clearance (50).

Our results support that CB2 plays a role in microglial response and phenotype. A previous study reported that CB2-deficient microglia display shorter projections that are less ramified within 24 hours after Asyn PFF injections relative to CB2 wild-type microglia as measured by a rigorous Scholl analysis (21). Although, we sampled adequately throughout the nigra, our data did not show statistically significant differences in the size of IBA1+ cell bodies when evaluated by IHC at 8-weeks, yet our flow cytometry data does suggest that microglia exhibited a stronger activation profile as shown by the elevated CD11b expression. Importantly, consistent with our findings using CB2 inverse agonist treatment, Feng and colleagues similarly identified upregulated CD68+ microglia from the genetic ablation of CB2. It has previously been shown that PD patient monocytes have impaired phagocytic capabilities and reduced ability to clear Asyn (51, 52), therefore modulation of CB2 may lead to a therapeutic gain-of-function involving immune cell uptake and clearance. Given that SMM-189 dampened microglial activation (as indicated by reduced CD11b expression) and delayed monocyte infiltration in the brain at earlier timepoints (4 weeks) and then increased activation in both myeloid and lymphoid compartments at later timepoints (8 weeks), we conclude that CB2 modulation can potentially lead to dynamic changes in the extent of microglial activation overtime.

Although our study suggests that the beneficial effects of SMM-189 were due to modulation of the neuroinflammatory response, the mechanism(s) underlying this critical therapeutic effect have not been fully elucidated. Previous work has identified CB2-mediated cAMP/PKA signaling pathways including the phosphorylation of CREB to impact the microglial phenotype (21, 26, 53). Additionally, recent reports show that microglial transcription of CB2 is dependent on nuclear factor erythroid 2-related factor 2 (NRF2), a well-known regulator of redox homeostasis and inflammation (54). An antioxidant response element (ARE) sequence was recently found in the promoter region of the microglial CB2 (54). NRF2 binds ARE in the nucleus to initiate transcription, suggesting that CB2 is linked to microglia-specific NFR2 inflammatory signaling. We were not able to test the specific signaling of SMM-189 responsible for the microglial phenotype transformations. However, in future studies it will be necessary to evaluate the mechanisms of microglial CB2 modulation in the presence of a synucleinopathy as they may be time-dependent in a progressive model such as the one used herein.

Our flow cytometry analyses revealed SMM-189 decreased the extent of infiltrating monocytes and CD4+ T cells at 4 weeks following human Asyn overexpression. Other groups have shown that viral-mediated overexpression of Asyn in the brain induces an influx of peripheral chemokine receptor 2 or CCR2+ immune cells into the degenerating area and that knocking out CCR2 or blocking monocyte infiltration is sufficient to protect against neuronal degeneration (55). In line with reduced peripheral immune cell infiltration, microglia isolated from SMM-189 treated mice were less activated at 4 weeks as measured by a reduced CD11b+ MFI. Together, these data suggest that SMM-189 can modify early microglial responses to insults such as Asyn overexpression. Interestingly, SMM-189 treatment led to a higher frequency of monocytes in the lesioned hemisphere at 8 weeks following human Asyn overexpression compared to vehicle treatment, suggesting SMM-189 delayed the resolution of monocyte infiltration in the lesioned hemisphere. Interesting, at this timepoint reduced CD163-immunoreactivity was found in the lesioned hemisphere by IHC. Expression of CD163 in the brain has been associated with border associated macrophages, which have been identified as the driving cells for immune cell recruitment and infiltration (56). Since we only evaluated CD163 expresssion at 8 weeks, it is possible that we missed a significant increase in this glycoprotein. Yet, the heightened activation of brain immune cells at 8 weeks, including more CD11b+ and CD172a+ expression on microglia and frequency of MHCII+ monocytes, is likely to have triggered the extravasation and increased presence of CD4+ helper cells and decreased CD8+ cytotoxic cells. The increased CD4 to CD8 ratio found in the brains of SMM-189-treated rats support an activated immune response that is likely to have contributed to the reduced synuclein pathology. Previous reports have found that T cell adoptive transfer to immune deficient mice reduced phosphorylated Asyn after pre-formed fibril injections, suggesting that T cells can have direct modulatory effects on Asyn toxicity (57). It is possible that CB2 modulation had direct effects on T cells as they also express the receptor and CB2-mediated activation of T cells alter functions including proliferation and migration (58–60). Although our results did show changes in T cell frequencies in the brain, the peripheral CD4+ and CD8+ T cells frequency were not altered from systemic SMM-189 treatment. Therefore, our data supports a model in which CB2-dependent modulation of brain myeloid cell function serves to attract and activate brain-infiltrating lymphocytes with little effect on peripheral T cells. We were not able to measure subtypes of T cells in the brain or periphery, with the exception of Tregs, a cell essential to regulating and suppressing immune cells. We found increased peripheral Tregs after 8 weeks of peripheral SMM-189 treatment compared to baseline (**SFigure 3**). Positive effects of Tregs in models of synucleiopathy have been reported by several groups (61, 62).

Peripheral treatment with CB2 inverse agonist SMM-189 produced body-wide effects that were independent of human Asyn overexpression. SMM-189 was administered peripherally and therefore is expected to elicit widespread effects in the brain as demonstrated by immune gene expression changes in the frontal cortex, an area not exposed directly human Asyn (**Figure 3H**). Furthermore, through multiple comparison analysis of the flow cytometry data, some of the immune effects were found in the non-lesioned hemisphere and not in the hemisphere overexpressing human Asyn. This is not surprising as intracerebral injections break the dura and could disturb the entire brain immune homeostasis, which in turn can affect immune cell infiltration throughout the brain, not just in the injected hemisphere. Our peripheral flow data at 8 weeks revealed that SMM-189 skewed peripheral monocytes with decreased pro-inflammatory CD43-His48+ classical monocytes and increased wound-healing CD43+ non-classical monocytes. Although neither specific subset of monocytes was significantly altered in the brain, the shift in microglial phenotypes may have increased the amount of CCL2 release leading to an overall elevated monocyte fraction in SMM-189 lesioned hemispheres. Additionally, there was a significant shift in peripheral immune phenotype based on PBMC gene expression, with decreased TNF and increased TGFβ which promotes alternative macrophage activation. These findings suggest that SMM-189 altered downstream CB2 signaling and promoted an anti-inflammatory brain microenvironment in the nigra that would not have been captured by evaluating biofluids at a systems level.

## Conclusion

The cannabinoid system has become a new and exciting target for immunomodulation in PD and other neurological conditions (20). Besides the recent genetic link to the cannabinoid system, others have identified elevated protein expression of CB2 in brains of individuals with PD, underscoring the neuroanatomical basis for targeting CB2. Our novel findings demonstrate that a CB2-selective inverse agonist, SMM-189, can alter immune responses both in the periphery and the brain to mitigate toxic species of Asyn in the brain. To our knowledge, this is the first study to highlight CB2 as a promising therapeutic target via which modulation of inflammatory and immune responses could delay, limit or prevent proteinopathies.

## List of abbreviations

CB2: cannabinoid receptor 2
Asyn: alpha-synuclein
pSer129: phosphorylated serine 129 alpha-synuclein
PD: Parkinson’s disease
LPS: lipopolysaccharide
PBMC: peripheral blood mononuclear cells
MFI: mean fluorescent intensity

## Declarations

### Ethics approval and consent to participate

Sprague Dawley rats were housed and handled in accordance with protocols approved by the IACUC of Emory University (protocol: DAR-2003358-ENTRPR-N). All surgeries and procedures including those under anesthesia were performed in IACUC approved settings at Emory University. Additionally, anesthesia and euthanasia procedures were conducted in accordance with the American Veterinary Medical Association Guidelines for the Euthanasia of animals and approved by Division of Animal Resources veterinarians at Emory. Analysis of the brain-to-blood ratios of SMM-189 were performed by Sai Life Sciences Limited (Telangana, India). All procedures of the study were in accordance with the guidelines provided by the Committee for the Purpose of Control and Supervision of Experiments on Animals (CPCSEA) as published in The Gazette of India, December 15, 1998. Prior approval of the Institutional Animal Ethics Committee (IAEC) was obtained before initiation of the study.

#### Consent for publication

Not applicable

#### Availability of data and materials

All data generated or analyzed during this study are included in this published article [and its supplementary information files]. Raw files can be provided upon request to the corresponding author.

#### Funding

Funding for this study was provided by the following: The Michael J Fox Foundation Target validation grant 2016 (MGT) and its supplement (VJ), Graduate and Postdoctoral Training in Environmental Health Science and Toxicology T32ES012870 (VJ), Emory Udall Pilot grant 1P50NS098685 (VJ), Training in translational Research in Neurology 2T32NS007480-16 (VJ). Funding for the murine PK study was provided by a seed grant from the University of Tennessee Health Science Center (BMM).

### Authors contributions

VJ and MGT conceived the study. VJ designed the study and supervised the technical staff involved in the study. VJ, FPM, BM, CM, DO, HS, JC, CAM, SG, SDK, MG, DL performed the various surgeries and assays. BMM provided guidance on drug dosing and study design. VJ, BMM, and MGT wrote and edited the paper. All authors reviewed and approved the final version of the manuscript.

## Acknowledgements

We thank the Emory Flow Cytometry Core and Emory Multiplexed Immunoassay Core (EMIC) for their assistance with the use of the LSRII and performing MSD multi-plex immunoassays, respectively. We thank the Tansey lab for thoughtful discussions.

## Supplemental Figures

**Table S1.**
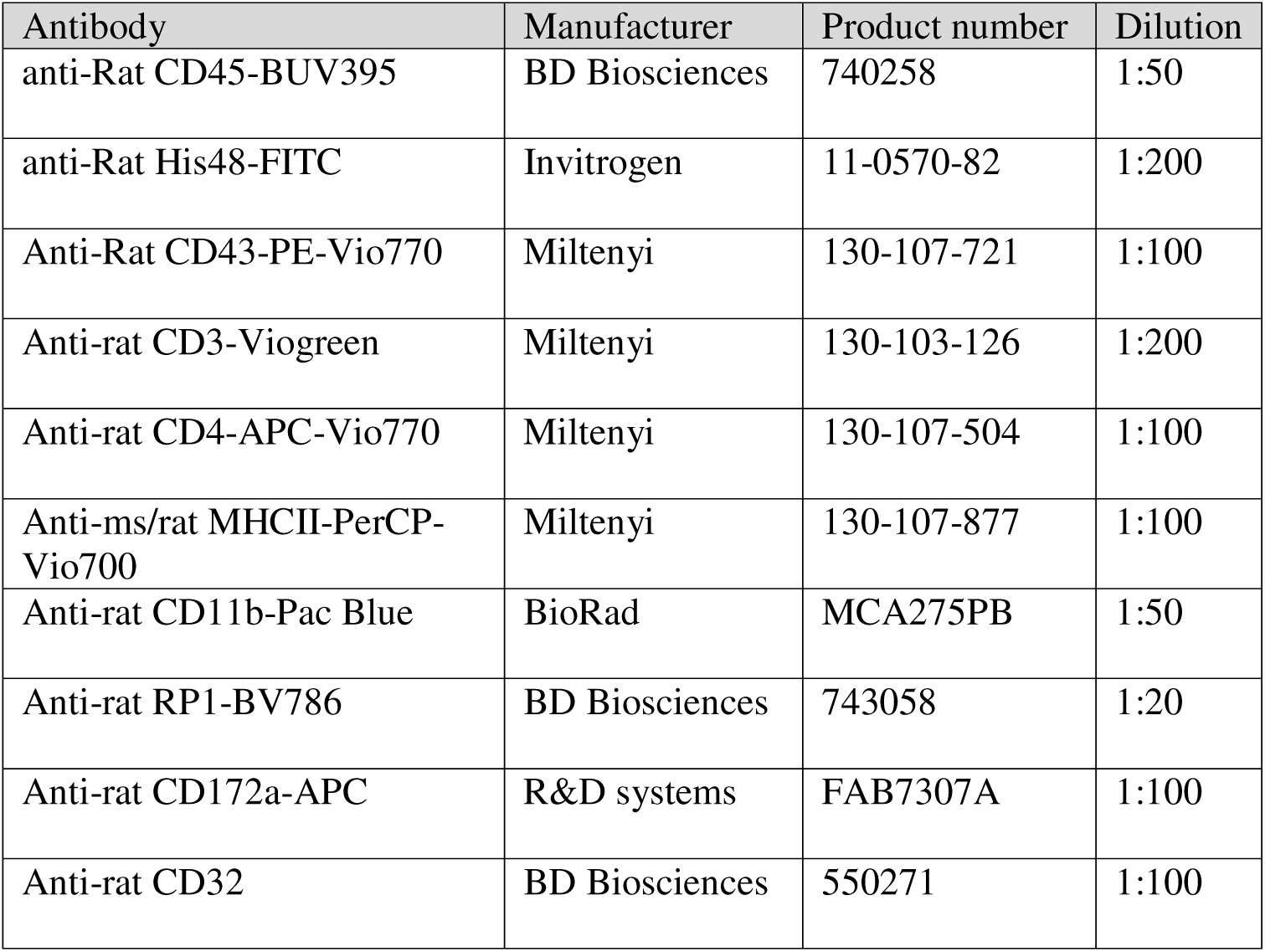
Myeloid PBMC flow panel.

**Table S2.** Lymphoid PBMC flow panel.

| Antibody | Manufacturer | Product number | Dilution |
| --- | --- | --- | --- |
| Anti-rat CD45RA-PECy7 | Biolegend | 202315 | 1:200 |
| Anti-rat CD3-Viogreen | Miltenyi | 130-103-126 | 1:200 |
| Anti-rat CD4-APC-Vio770 | Miltenyi | 130-107-504 | 1:100 |
| Anti-CD8a-PerCP-Vio700 | Miltenyi | 130-108-914 | 1:100 |
| Anti-rat CD25-BV786 | BD Biosciences | 742757 | 1:50 |
| Anti-ms/rat Foxp3-AF647<br>*stained following<br>intracellular permeabilization | R&D systems | FAB7307A | 1:50 |
| Anti-rat CD32 | BD Biosciences | 550271 | 1:100 |

**Table S3.** Brain Immune cell flow panel.

| Antibody | Manufacturer | Product number | Dilution |
| --- | --- | --- | --- |
| anti-Rat CD44H-FITC | Miltenyi | 130-107-854 | 1:50 |
| Anti-Rat CD62L-PE | BD Biosciences | 551398 | 1:200 |
| Anti-rat CD45RA-PECy7 | Biolegend | 202315 | 1:200 |
| Anti-rat CD3-Viogreen | Miltenyi | 130-103-126 | 1:200 |
| Anti-rat CD4-APC-Vio770 | Miltenyi | 130-107-504 | 1:100 |
| Anti-CD8a-PerCP-Vio700 | Miltenyi | 130-108-914 | 1:100 |
| Anti-rat CD25-BV786 | BD Biosciences | 742757 | 1:50 |
| Anti-ms/rat Foxp3-AF647<br>*stained following<br>intracellular permeabilization | R&D systems | FAB7307A | 1:50 |
| Anti-rat CD32 | BD Biosciences | 550271 | 1:100 |

**Table S4.**
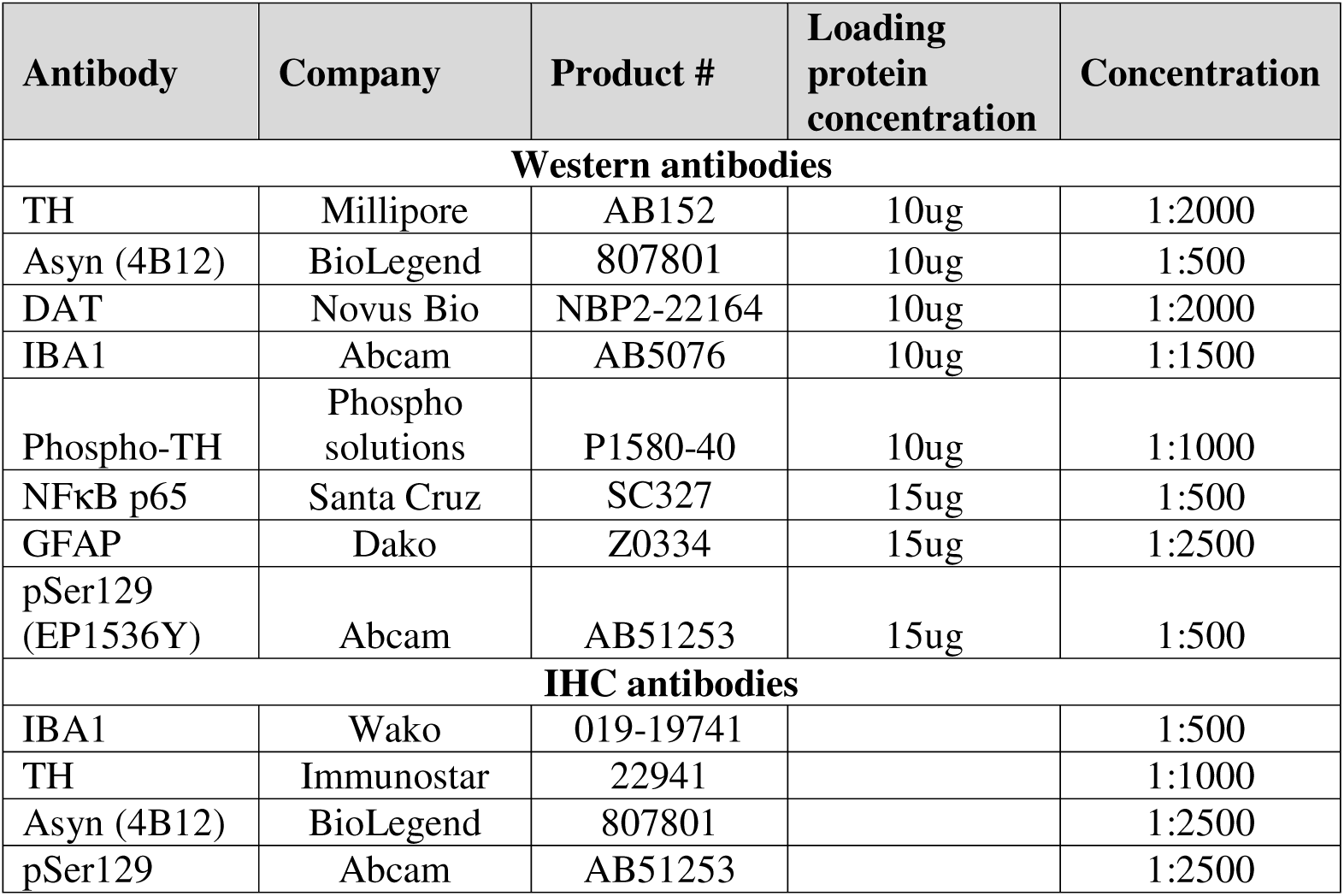
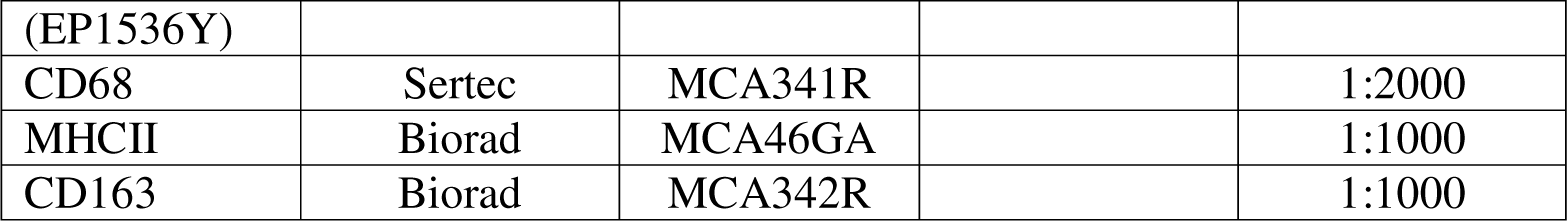
Western and IHC antibodies.

## References

1. Uriarte Huarte O, Kyriakis D, Heurtaux T, Pires-Afonso Y, Grzyb K, Halder R, et al. Single-Cell Transcriptomics and In Situ Morphological Analyses Reveal Microglia Heterogeneity Across the Nigrostriatal Pathway. Front Immunol. 2021;12:639613.

2. Paolicelli RC, Sierra A, Stevens B, Tremblay ME, Aguzzi A, Ajami B, et al. Microglia states and nomenclature: A field at its crossroads. Neuron. 2022;110(21):3458–83.

3. Smajic S, Prada-Medina CA, Landoulsi Z, Ghelfi J, Delcambre S, Dietrich C, et al. Single-cell sequencing of human midbrain reveals glial activation and a Parkinson-specific neuronal state. Brain. 2022;145(3):964–78.

4. Mastroeni D, Nolz J, Sekar S, Delvaux E, Serrano G, Cuyugan L, et al. Laser-captured microglia in the Alzheimer’s and Parkinson’s brain reveal unique regional expression profiles and suggest a potential role for hepatitis B in the Alzheimer’s brain. Neurobiol Aging. 2018;63:12–21.

5. Joers V, Tansey MG, Mulas G, Carta AR. Microglial phenotypes in Parkinson’s disease and animal models of the disease. Prog Neurobiol. 2017;155:57–75.

6. McGeer PL, Itagaki S, Boyes BE, McGeer EG. Reactive microglia are positive for HLA-DR in the substantia nigra of Parkinson’s and Alzheimer’s disease brains. Neurology. 1988;38(8):1285–91.

7. Ferreira SA, Romero-Ramos M. Microglia Response During Parkinson’s Disease: Alpha-Synuclein Intervention. Front Cell Neurosci. 2018;12:247.

8. Earls RH, Menees KB, Chung J, Barber J, Gutekunst CA, Hazim MG, et al. Intrastriatal injection of preformed alpha-synuclein fibrils alters central and peripheral immune cell profiles in non-transgenic mice. J Neuroinflammation. 2019;16(1):250.

9. Theodore S, Cao S, McLean PJ, Standaert DG. Targeted overexpression of human alpha-synuclein triggers microglial activation and an adaptive immune response in a mouse model of Parkinson disease. J Neuropathol Exp Neurol. 2008;67(12):1149–58.

10. Gubinelli F, Sarauskyte L, Venuti C, Kulacz I, Cazzolla G, Negrini M, et al. Characterisation of functional deficits induced by AAV overexpression of alpha-synuclein in rats. Curr Res Neurobiol. 2023;4:100065.

11. Liu Z, Yang N, Dong J, Tian W, Chang L, Ma J, et al. Deficiency in endocannabinoid synthase DAGLB contributes to early onset Parkinsonism and murine nigral dopaminergic neuron dysfunction. Nat Commun. 2022;13(1):3490.

12. Ashton JC, Glass M. The cannabinoid CB2 receptor as a target for inflammation-dependent neurodegeneration. Curr Neuropharmacol. 2007;5(2):73–80.

13. Gomez-Galvez Y, Palomo-Garo C, Fernandez-Ruiz J, Garcia C. Potential of the cannabinoid CB(2) receptor as a pharmacological target against inflammation in Parkinson’s disease. Prog Neuropsychopharmacol Biol Psychiatry. 2016;64:200–8.

14. Garcia MC, Cinquina V, Palomo-Garo C, Rabano A, Fernandez-Ruiz J. Identification of CB(2) receptors in human nigral neurons that degenerate in Parkinson’s disease. Neurosci Lett. 2015;587:1–4.

15. Navarrete F, Garcia-Gutierrez MS, Aracil-Fernandez A, Lanciego JL, Manzanares J. Cannabinoid CB1 and CB2 Receptors, and Monoacylglycerol Lipase Gene Expression Alterations in the Basal Ganglia of Patients with Parkinson’s Disease. Neurotherapeutics. 2018;15(2):459–69.

16. Price DA, Martinez AA, Seillier A, Koek W, Acosta Y, Fernandez E, et al. WIN55,212-2, a cannabinoid receptor agonist, protects against nigrostriatal cell loss in the 1-methyl-4-phenyl-1,2,3,6-tetrahydropyridine mouse model of Parkinson’s disease. Eur J Neurosci. 2009;29(11):2177–86.

17. Garcia C, Palomo-Garo C, Garcia-Arencibia M, Ramos J, Pertwee R, Fernandez-Ruiz J. Symptom-relieving and neuroprotective effects of the phytocannabinoid Delta(9)-THCV in animal models of Parkinson’s disease. Br J Pharmacol. 2011;163(7):1495–506.

18. Ternianov A, Perez-Ortiz JM, Solesio ME, Garcia-Gutierrez MS, Ortega-Alvaro A, Navarrete F, et al. Overexpression of CB2 cannabinoid receptors results in neuroprotection against behavioral and neurochemical alterations induced by intracaudate administration of 6-hydroxydopamine. Neurobiol Aging. 2012;33(2):421 e1-16.

19. Javed H, Azimullah S, Haque ME, Ojha SK. Cannabinoid Type 2 (CB2) Receptors Activation Protects against Oxidative Stress and Neuroinflammation Associated Dopaminergic Neurodegeneration in Rotenone Model of Parkinson’s Disease. Front Neurosci. 2016;10:321.

20. Kelly R, Joers V, Tansey MG, McKernan DP, Dowd E. Microglial Phenotypes and Their Relationship to the Cannabinoid System: Therapeutic Implications for Parkinson’s Disease. Molecules. 2020;25(3).

21. Feng L, Lo H, You H, Wu W, Cheng X, Xin J, et al. Loss of cannabinoid receptor 2 promotes alpha-Synuclein-induced microglial synaptic pruning in nucleus accumbens by modulating the pCREB-c-Fos signaling pathway and complement system. Exp Neurol. 2023;359:114230.

22. Reiner A, Heldt SA, Presley CS, Guley NH, Elberger AJ, Deng Y, et al. Motor, visual and emotional deficits in mice after closed-head mild traumatic brain injury are alleviated by the novel CB2 inverse agonist SMM-189. Int J Mol Sci. 2014;16(1):758–87.

23. Presley C, Abidi A, Suryawanshi S, Mustafa S, Meibohm B, Moore BM. Preclinical evaluation of SMM-189, a cannabinoid receptor 2-specific inverse agonist. Pharmacol Res Perspect. 2015;3(4):e00159.

24. Guley NM, Del Mar NA, Ragsdale T, Li C, Perry AM, Moore BM, et al. Amelioration of visual deficits and visual system pathology after mild TBI with the cannabinoid type-2 receptor inverse agonist SMM-189. Exp Eye Res. 2019;182:109–24.

25. Atwood BK, Wager-Miller J, Haskins C, Straiker A, Mackie K. Functional selectivity in CB(2) cannabinoid receptor signaling and regulation: implications for the therapeutic potential of CB(2) ligands. Mol Pharmacol. 2012;81(2):250–63.

26. Bu W, Ren H, Deng Y, Del Mar N, Guley NM, Moore BM, et al. Mild Traumatic Brain Injury Produces Neuron Loss That Can Be Rescued by Modulating Microglial Activation Using a CB2 Receptor Inverse Agonist. Front Neurosci. 2016;10:449.

27. Gombash SE, Manfredsson FP, Kemp CJ, Kuhn NC, Fleming SM, Egan AE, et al. Morphological and behavioral impact of AAV2/5-mediated overexpression of human wildtype alpha-synuclein in the rat nigrostriatal system. PLoS One. 2013;8(11):e81426.

28. Sandoval IM, Kuhn NM, Manfredsson FP. Multimodal Production of Adeno-Associated Virus. Methods Mol Biol. 2019;1937:101–24.

29. Benskey MJ, Sandoval IM, Manfredsson FP. Continuous Collection of Adeno-Associated Virus from Producer Cell Medium Significantly Increases Total Viral Yield. Hum Gene Ther Methods. 2016;27(1):32–45.

30. Benskey MJ, Manfredsson FP. Intraparenchymal Stereotaxic Delivery of rAAV and Special Considerations in Vector Handling. Methods Mol Biol. 2016;1382:199–215.

31. Alghamdi SS, Mustafa SM, Moore Ii BM. Synthesis and biological evaluation of a ring analogs of the selective CB2 inverse agonist SMM-189. Bioorg Med Chem. 2021;33:116035.

32. Houser MC, Uriarte Huarte O, Wallings RL, Keating CE, MacPherson KP, Herrick MK, et al. Progranulin loss results in sex-dependent dysregulation of the peripheral and central immune system. Front Immunol. 2022;13:1056417.

33. MacPherson KP, Eidson LN, Houser MC, Weiss BE, Gollihue JL, Herrick MK, et al. Soluble TNF mediates amyloid-independent, diet-induced alterations to immune and neuronal functions in an Alzheimer’s disease mouse model. Front Cell Neurosci. 2023;17:895017.

34. Barnett-Vanes A, Sharrock A, Birrell MA, Rankin S. A Single 9-Colour Flow Cytometric Method to Characterise Major Leukocyte Populations in the Rat: Validation in a Model of LPS-Induced Pulmonary Inflammation. PLoS One. 2016;11(1):e0142520.

35. Ahuja V, Miller SE, Howell DN. Identification of two subpopulations of rat monocytes expressing disparate molecular forms and quantities of CD43. Cell Immunol. 1995;163(1):59–69.

36. Kiefer J, Zeller J, Bogner B, Horbrand IA, Lang F, Deiss E, et al. An Unbiased Flow Cytometry-Based Approach to Assess Subset-Specific Circulating Monocyte Activation and Cytokine Profile in Whole Blood. Front Immunol. 2021;12:641224.

37. Houser MC, Caudle WM, Chang J, Kannarkat GT, Yang Y, Kelly SD, et al. Experimental colitis promotes sustained, sex-dependent, T-cell-associated neuroinflammation and parkinsonian neuropathology. Acta Neuropathol Commun. 2021;9(1):139.

38. Joers V, Masilamoni G, Kempf D, Weiss AR, Rotterman TM, Murray B, et al. Microglia, inflammation and gut microbiota responses in a progressive monkey model of Parkinson’s disease: A case series. Neurobiol Dis. 2020;144:105027.

39. Fujiwara H, Hasegawa M, Dohmae N, Kawashima A, Masliah E, Goldberg MS, et al. alpha-Synuclein is phosphorylated in synucleinopathy lesions. Nat Cell Biol. 2002;4(2):160–4.

40. Anderson JP, Walker DE, Goldstein JM, de Laat R, Banducci K, Caccavello RJ, et al. Phosphorylation of Ser-129 is the dominant pathological modification of alpha-synuclein in familial and sporadic Lewy body disease. J Biol Chem. 2006;281(40):29739–52.

41. Smith WW, Margolis RL, Li X, Troncoso JC, Lee MK, Dawson VL, et al. Alpha-synuclein phosphorylation enhances eosinophilic cytoplasmic inclusion formation in SH-SY5Y cells. J Neurosci. 2005;25(23):5544–52.

42. Karampetsou M, Ardah MT, Semitekolou M, Polissidis A, Samiotaki M, Kalomoiri M, et al. Phosphorylated exogenous alpha-synuclein fibrils exacerbate pathology and induce neuronal dysfunction in mice. Sci Rep. 2017;7(1):16533.

43. Tansey MG, Wallings RL, Houser MC, Herrick MK, Keating CE, Joers V. Inflammation and immune dysfunction in Parkinson disease. Nat Rev Immunol. 2022;22(11):657–73.

44. Lopez A, Aparicio N, Pazos MR, Grande MT, Barreda-Manso MA, Benito-Cuesta I, et al. Cannabinoid CB(2) receptors in the mouse brain: relevance for Alzheimer’s disease. J Neuroinflammation. 2018;15(1):158.

45. Schmole AC, Lundt R, Toporowski G, Hansen JN, Beins E, Halle A, et al. Cannabinoid Receptor 2-Deficiency Ameliorates Disease Symptoms in a Mouse Model with Alzheimer’s Disease-Like Pathology. J Alzheimers Dis. 2018;64(2):379–92.

46. Galan-Ganga M, Rodriguez-Cueto C, Merchan-Rubira J, Hernandez F, Avila J, Posada-Ayala M, et al. Cannabinoid receptor CB2 ablation protects against TAU induced neurodegeneration. Acta Neuropathol Commun. 2021;9(1):90.

47. Danzer KM, Kranich LR, Ruf WP, Cagsal-Getkin O, Winslow AR, Zhu L, et al. Exosomal cell-to-cell transmission of alpha synuclein oligomers. Mol Neurodegener. 2012;7:42.

48. Desplats P, Lee HJ, Bae EJ, Patrick C, Rockenstein E, Crews L, et al. Inclusion formation and neuronal cell death through neuron-to-neuron transmission of alpha-synuclein. Proc Natl Acad Sci U S A. 2009;106(31):13010–5.

49. Lee KW, Chen W, Junn E, Im JY, Grosso H, Sonsalla PK, et al. Enhanced phosphatase activity attenuates alpha-synucleinopathy in a mouse model. J Neurosci. 2011;31(19):6963–71.

50. Barko K, Shelton M, Xue X, Afriyie-Agyemang Y, Puig S, Freyberg Z, et al. Brain region- and sex-specific transcriptional profiles of microglia. Front Psychiatry. 2022;13:945548.

51. Bliederhaeuser C, Grozdanov V, Speidel A, Zondler L, Ruf WP, Bayer H, et al. Age-dependent defects of alpha-synuclein oligomer uptake in microglia and monocytes. Acta Neuropathol. 2016;131(3):379–91.

52. Grozdanov V, Bliederhaeuser C, Ruf WP, Roth V, Fundel-Clemens K, Zondler L, et al. Inflammatory dysregulation of blood monocytes in Parkinson’s disease patients. Acta Neuropathol. 2014;128(5):651–63.

53. Tao Y, Li L, Jiang B, Feng Z, Yang L, Tang J, et al. Cannabinoid receptor-2 stimulation suppresses neuroinflammation by regulating microglial M1/M2 polarization through the cAMP/PKA pathway in an experimental GMH rat model. Brain Behav Immun. 2016;58:118–29.

54. Galan-Ganga M, Del Rio R, Jimenez-Moreno N, Diaz-Guerra M, Lastres-Becker I. Cannabinoid CB(2) Receptor Modulation by the Transcription Factor NRF2 is Specific in Microglial Cells. Cell Mol Neurobiol. 2020;40(1):167–77.

55. Harms AS, Thome AD, Yan Z, Schonhoff AM, Williams GP, Li X, et al. Peripheral monocyte entry is required for alpha-Synuclein induced inflammation and Neurodegeneration in a model of Parkinson disease. Exp Neurol. 2018;300:179–87.

56. Schonhoff AM, Figge DA, Williams GP, Jurkuvenaite A, Gallups NJ, Childers GM, et al. Border-associated macrophages mediate the neuroinflammatory response in an alpha-synuclein model of Parkinson disease. Nat Commun. 2023;14(1):3754.

57. George S, Tyson T, Rey NL, Sheridan R, Peelaerts W, Becker K, et al. T Cells Limit Accumulation of Aggregate Pathology Following Intrastriatal Injection of alpha-Synuclein Fibrils. J Parkinsons Dis. 2021;11(2):585–603.

58. Bouaboula M, Rinaldi M, Carayon P, Carillon C, Delpech B, Shire D, et al. Cannabinoid-receptor expression in human leukocytes. Eur J Biochem. 1993;214(1):173–80.

59. Castaneda JT, Harui A, Kiertscher SM, Roth JD, Roth MD. Differential expression of intracellular and extracellular CB(2) cannabinoid receptor protein by human peripheral blood leukocytes. J Neuroimmune Pharmacol. 2013;8(1):323–32.

60. Simard M, Rakotoarivelo V, Di Marzo V, Flamand N. Expression and Functions of the CB(2) Receptor in Human Leukocytes. Front Pharmacol. 2022;13:826400.

61. Badr M, McFleder RL, Wu J, Knorr S, Koprich JB, Hunig T, et al. Expansion of regulatory T cells by CD28 superagonistic antibodies attenuates neurodegeneration in A53T-alpha-synuclein Parkinson’s disease mice. J Neuroinflammation. 2022;19(1):319.

62. Sanchez-Guajardo V, Annibali A, Jensen PH, Romero-Ramos M. alpha-Synuclein vaccination prevents the accumulation of parkinson disease-like pathologic inclusions in striatum in association with regulatory T cell recruitment in a rat model. J Neuropathol Exp Neurol. 2013;72(7):624–45.

