## Supplemental figures for "Modulation of cannabinoid receptor 2 alters neuroinflammation and reduces formation of alpha-synuclein aggregates in a rat model of nigral synucleinopathy"

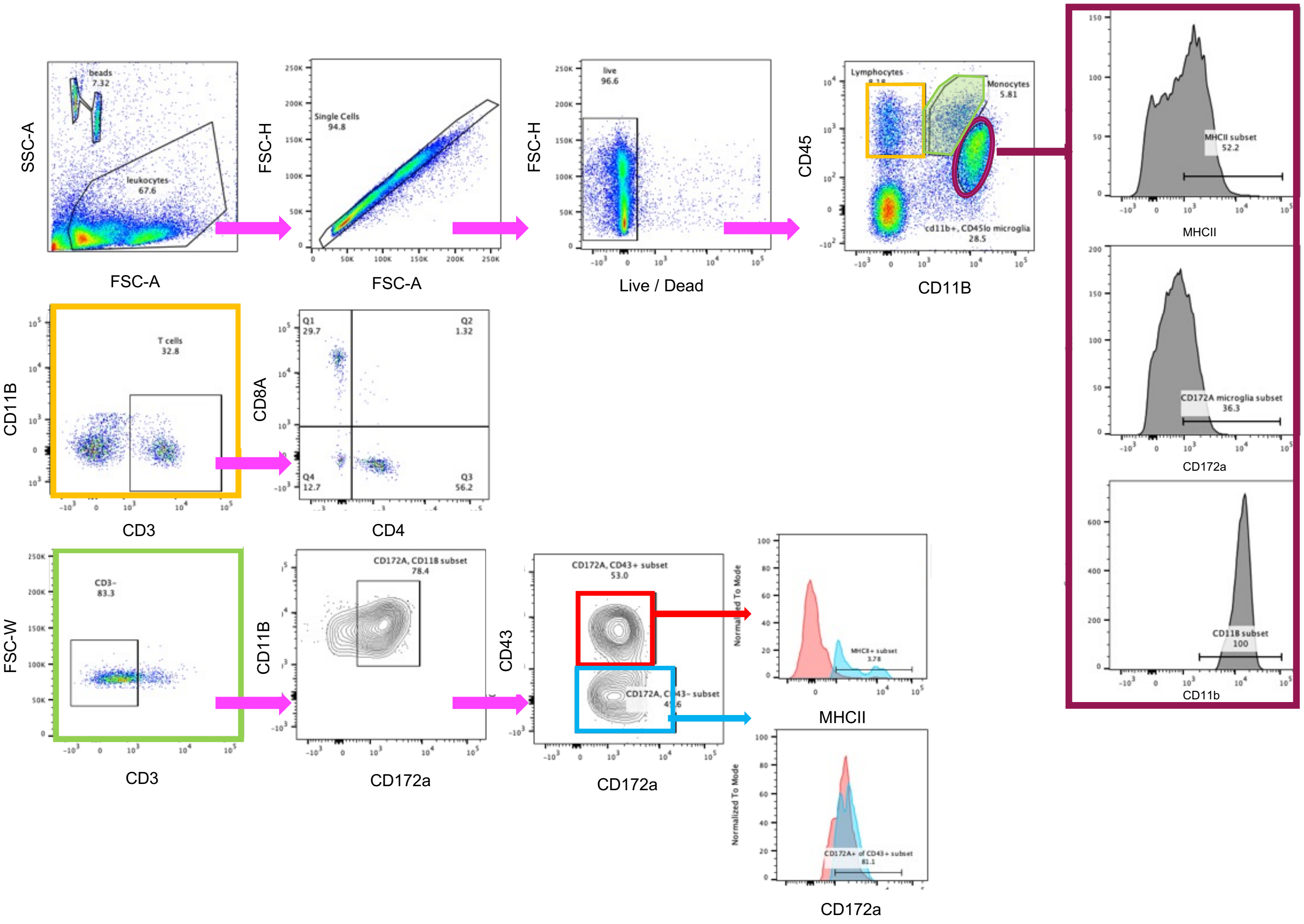


**Figure S1.** ***Flow gating strategy for rat brain immune cells***

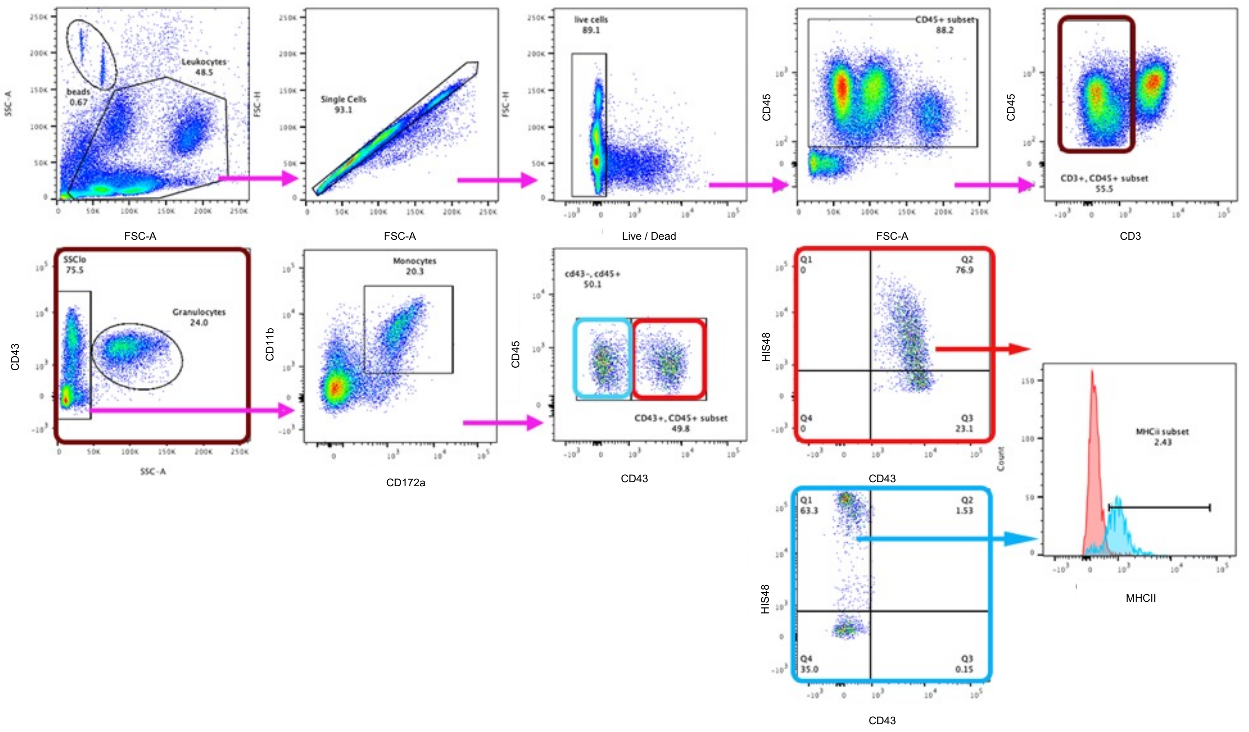


**Figure S2.** ***Flow gating strategy for rat PBMC myeloid cells***

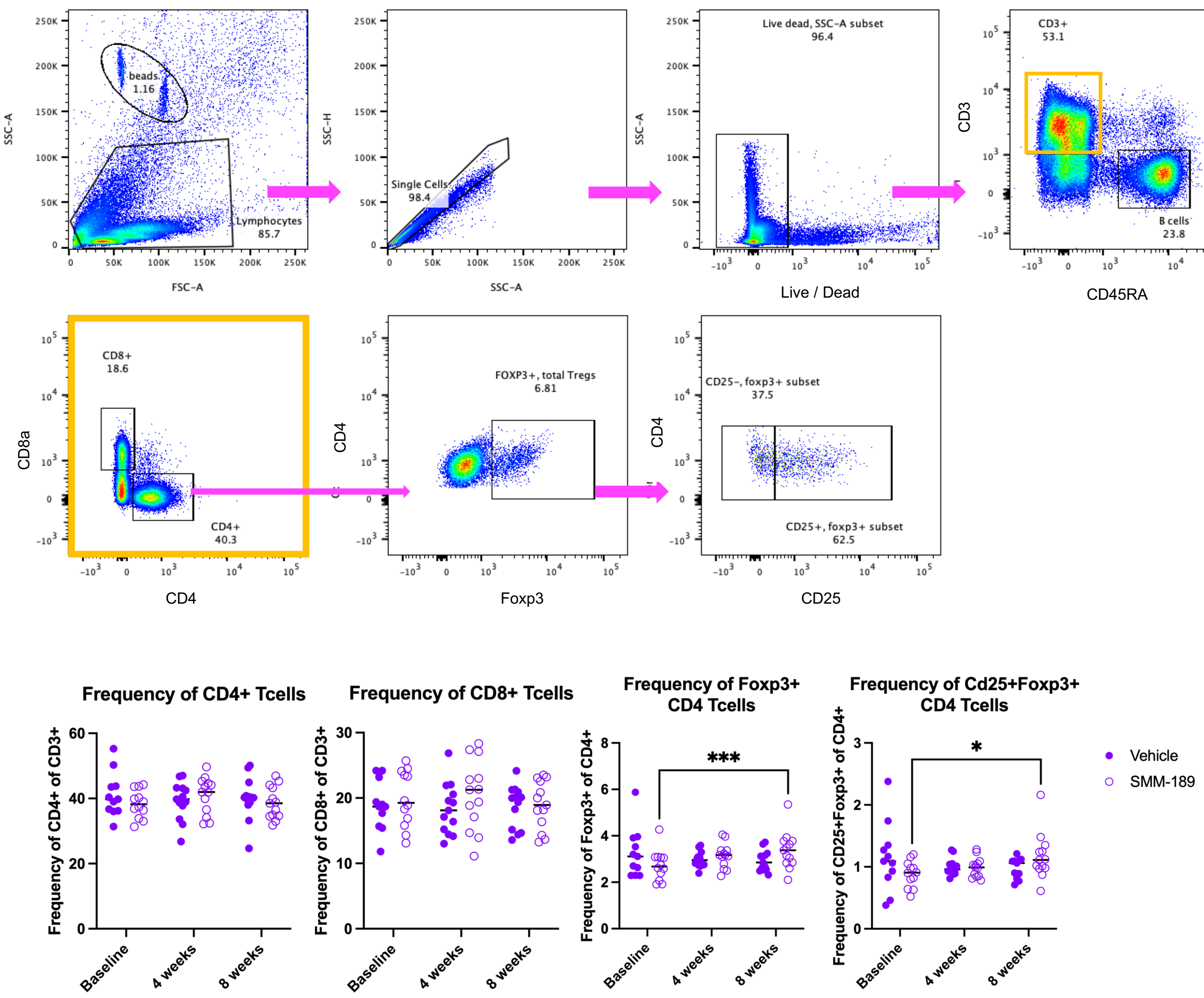


***Figure S3. Flow gating strategy for rat lymphocytes and elevation of peripheral Tregs in SMM-189-treated rats (high cohort) after 7 weeks of treatment***

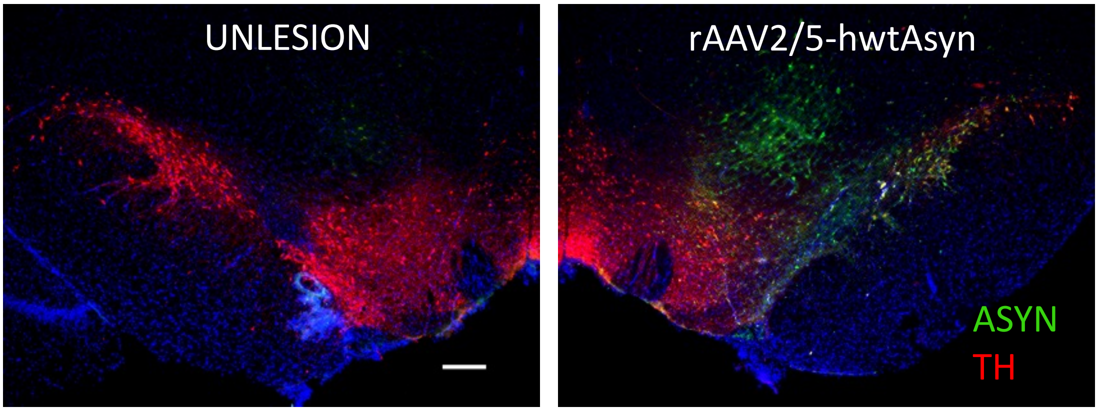


***Figure S4. Targeting of rAAV2/5-hwtAsyn in the rat substantia nigra***

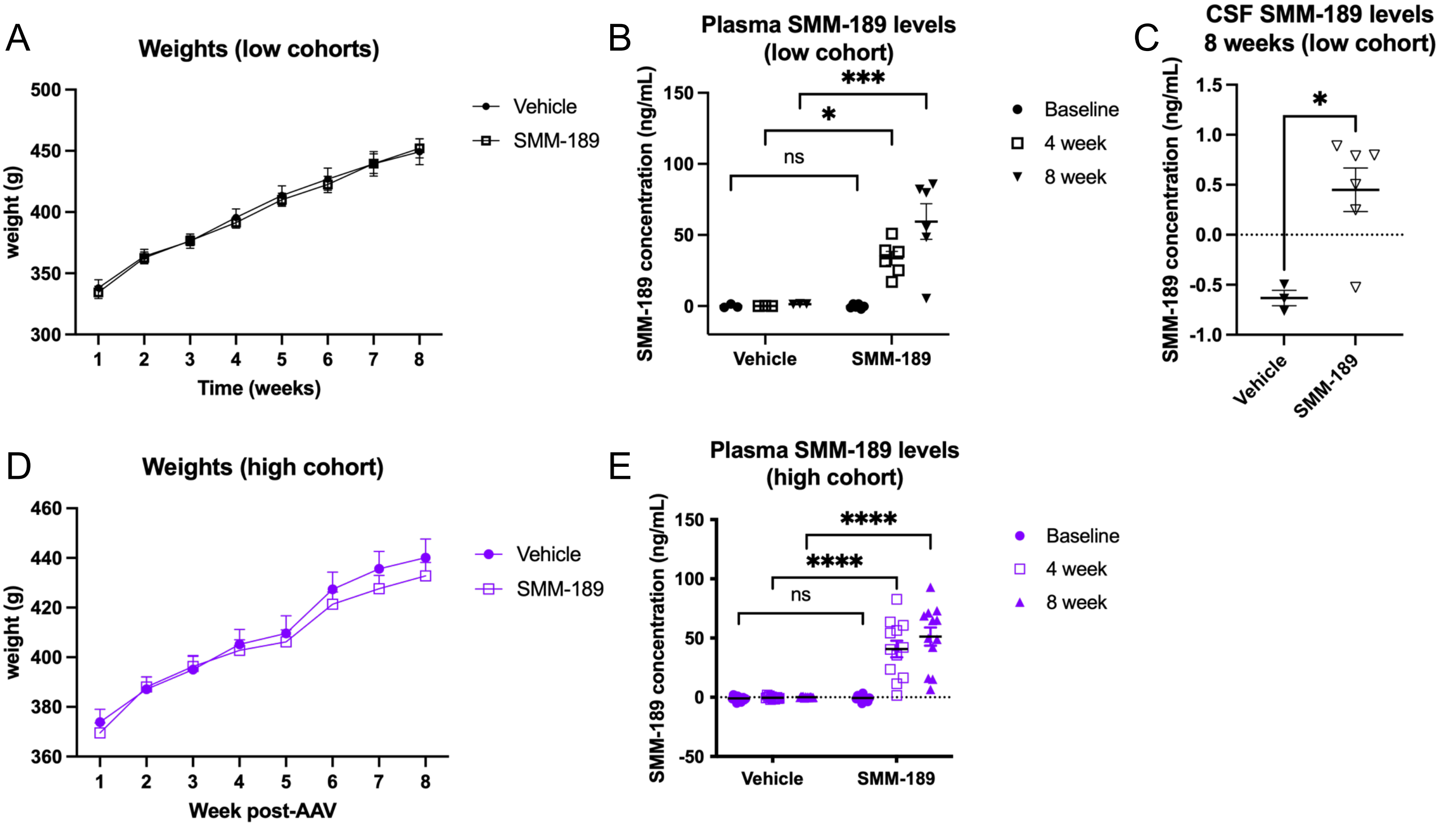


***Figure S5. Animal body weights and levels of SMM-189 in plasma and CSF***

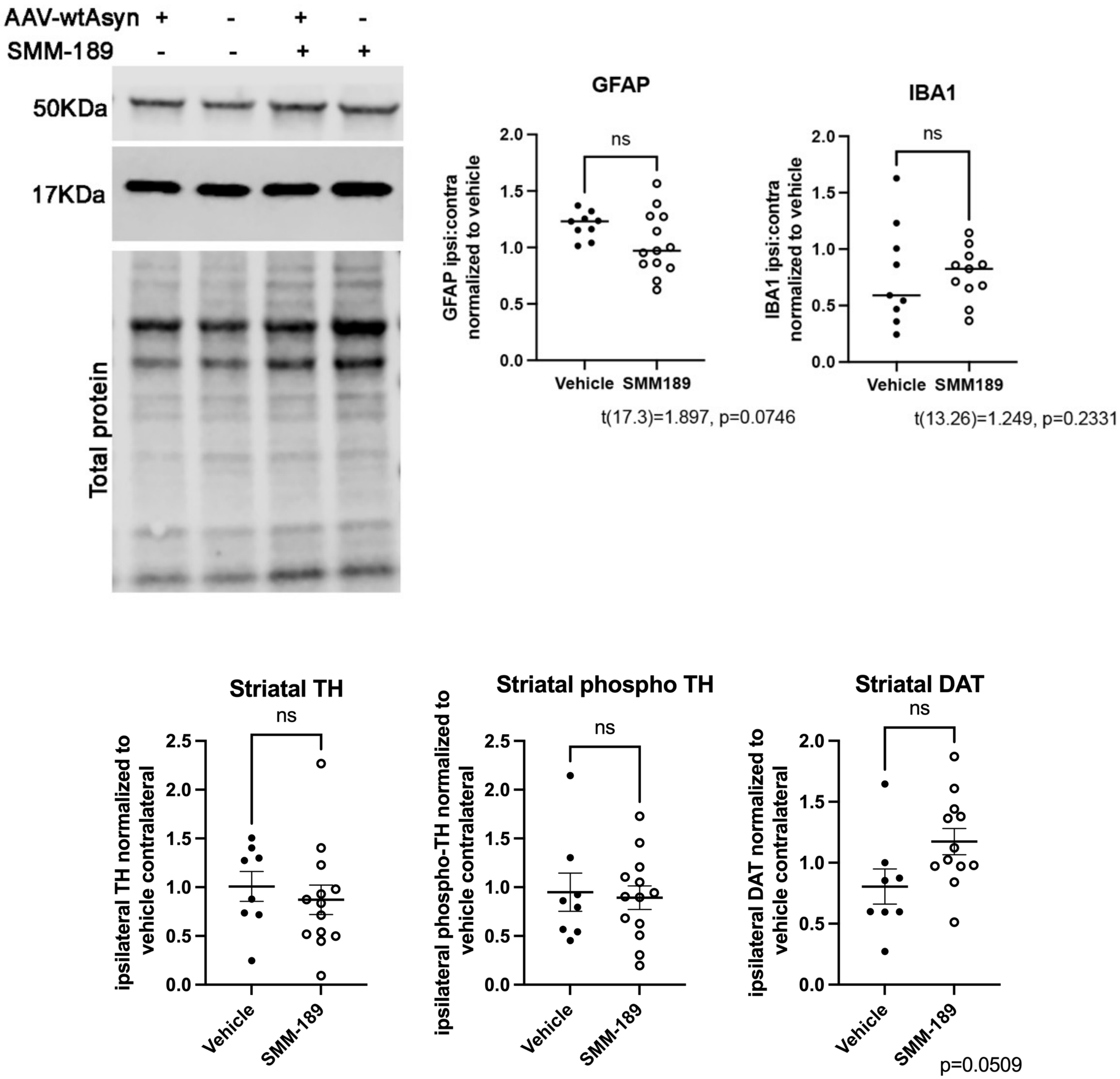


***Figure S6. Protein markers for astrocytes, microglia or dopaminergic neurons in striatum do not change by CB2 modulation with SMM-189***

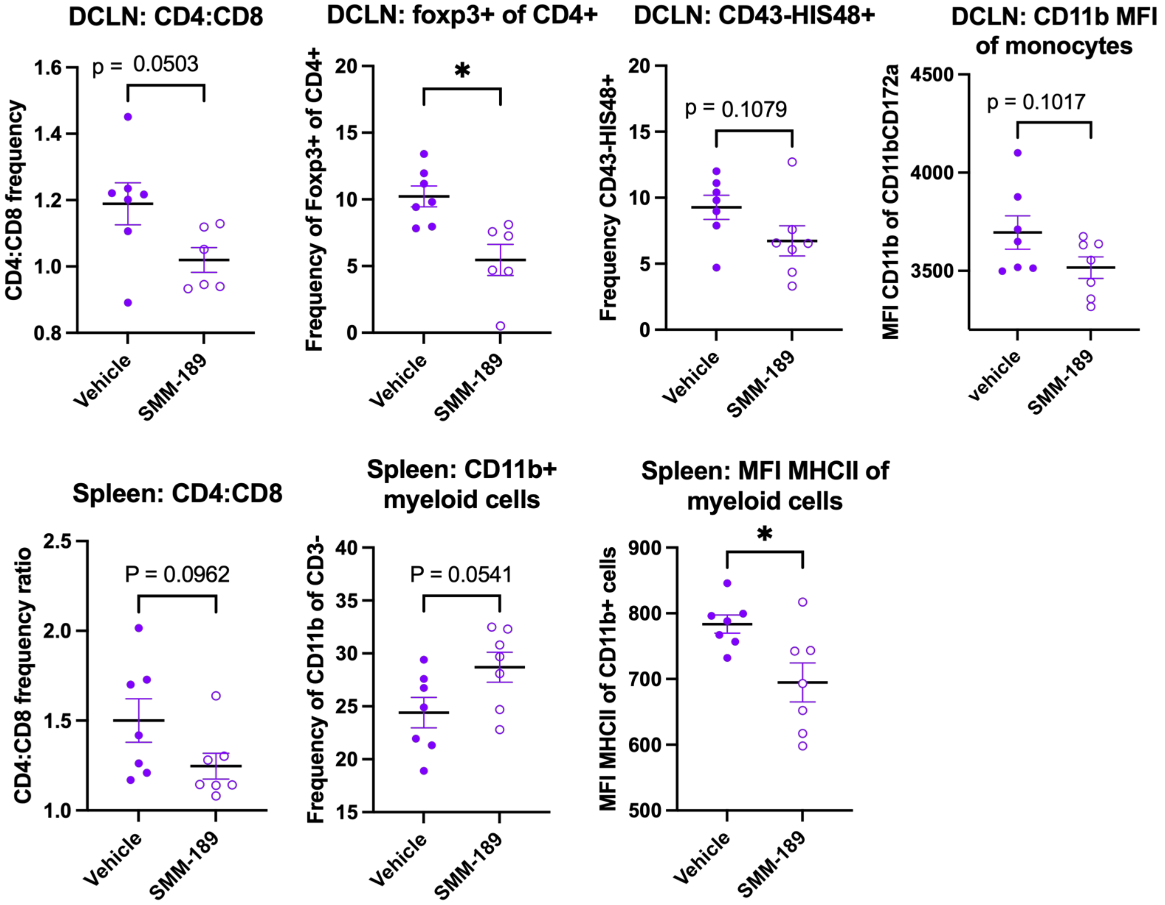


***Figure S7. Immune cell populations in deep-cervical lymph nodes (DCLN) and splenocytes are minimally affected in SMM-189-treated rats (high cohort) after 7 weeks of dosing***
